# Direct and indirect consequences of *PAB1* deletion in the regulation of translation initiation, translation termination, and mRNA decay

**DOI:** 10.1101/2023.05.31.543082

**Authors:** Kotchaphorn Mangkalaphiban, Robin Ganesan, Allan Jacobson

## Abstract

Cytoplasmic poly(A)-binding protein (PABPC; Pab1 in yeast) is thought to be involved in multiple steps of post-transcriptional control, including translation initiation, translation termination, and mRNA decay. To understand these roles of PABPC in more detail for endogenous mRNAs, and to distinguish its direct effects from indirect effects, we have employed RNA-Seq and Ribo-Seq to analyze changes in the abundance and translation of the yeast transcriptome, as well as mass spectrometry to assess the abundance of the components of the yeast proteome, in cells lacking the *PAB1* gene. We observed drastic changes in the transcriptome and proteome, as well as defects in translation initiation and termination, in *pab1Δ* cells. Defects in translation initiation and the stabilization of specific classes of mRNAs in *pab1Δ* cells appear to be partly indirect consequences of reduced levels of specific initiation factors, decapping activators, and components of the deadenylation complex in addition to the general loss of Pab1’s direct role in these processes. Cells devoid of Pab1 also manifested a nonsense codon readthrough phenotype indicative of a defect in translation termination, but this defect may be a direct effect of the loss of Pab1 as it could not be attributed to significant reductions in the levels of release factors.

**AUTHOR SUMMARY:** Many human diseases are caused by having too much or too little of certain cellular proteins. The amount of an individual protein is influenced by the level of its messenger mRNA (mRNA) and the efficiency of translation of the mRNA into a polypeptide chain by the ribosomes. Cytoplasmic poly(A)-binding protein (PABPC) plays numerous roles in the regulation of this multi-staged process, but understanding its specific role has been challenging because it is sometimes unclear whether experimental results are related to PABPC’s direct role in a specific biochemical process or to indirect effects of its other roles, leading to conflicting models of PABPC’s functions between studies. In this study, we characterized defects of each stage of protein synthesis in response to loss of PABPC in yeast cells by measuring whole-cell levels of mRNAs, ribosome-associated mRNAs, and proteins. We demonstrated that defects in most steps of protein synthesis other than the last can be explained by reduced levels of mRNAs that code for proteins important for that step in addition to loss of PABPC’s direct role on that step. Our data and analyses serve as resources for the design of future studies of PABPC’s functions.

## INTRODUCTION

Eukaryotic mRNAs are subject to complex post-transcriptional regulation by RNA binding proteins (RBPs) that control protein output. Cytoplasmic poly(A)-binding proteins (PABPCs) are RBPs that bind polyadenylated tails at mRNA 3’-ends and subsequently play roles in multiple stages of cytoplasmic mRNA regulation, from translation initiation to termination and mRNA decay [1–3]. The numerous functions of PABPC are attributed to its ability to interact with not only mRNAs but also various other proteins to form messenger ribonucleoprotein (mRNP) complexes [2]. PABPC’s conserved structure consists of three regions: i) four RNA recognition motif (RRM) domains, two of which (RRM1 and RRM2) bind 12 adenosines with high affinity [3,4], while the protein overall covers 27 adenosines [5], ii) a proline-rich (P) linker domain, and iii) a C-terminal mademoiselle (MLLE) domain that interacts with other proteins via their PABP-interacting motif 2 (PAM2) [2,3,6]. Mammals have multiple isoforms of PABPCs, of which the most ubiquitous isoform as well as the most studied is PABPC1, whereas the yeast *Saccharomyces cerevisiae* has only one PABPC, Pab1 [2,3,6,7].

The current model for general mRNA decay in yeast involves biphasic poly(A) tailing shortening by Pan2-Pan3 and Ccr4-Not deadenylase complexes, decapping by the Dcp1/Dcp2 holoenzyme, and exonucleolytic Xrn1-mediated 5’-3’ degradation or exosome-mediated 3’-5’ degradation [3,8,9]. PABPC appears to have a paradoxical role in these processes. On the one hand, PABPC1/Pab1 stimulates deadenylation by recruiting Pan2-Pan3 to the poly(A) tail through its interaction with the PAM2 motif on Pan3 [1,3,8,10,11]. Consistent with this model, mRNAs in yeast cells harboring a deletion of *PAB1* or mutations of Pab1’s C-terminal Pan3-interacting domain had longer poly(A) tails than their counterparts in wild-type cells [1–3,12–14]. On the other hand, PABPC has been shown to protect mRNAs from exonucleases [3,8].

Importantly, mRNAs with longer poly(A) tails are generally more stable and better translated than their short-tailed or un-tailed counterparts [3,15–23], observations that led us to propose a role for PABPC in the enhancement of translation initiation by the possible formation of an mRNA “closed-loop” [23–25]. Current elaborations of this model postulate that mRNAs could be circularized by a chain of interactions between poly(A)-associated PABPC and 5’ end-localized initiation factors, giving rise to a poly(A) tail-PABPC-eIF4G eIF4E-5’cap network [2,26–30]. eIF4G has been shown to interact with PABPC’s RRM2 domain [26,31–33], and disrupting this interaction reduced translation [6,29,34]. Thus, PABPC can stabilize the cap-binding complex and aid the recruitment of the 43S pre initiation complex to the mRNA [2,3,35]. However, this closed-loop arrangement is not required for all mRNAs or all conditions, raising the question of how else 5’-3’ communication is facilitated or whether it is a universal step [30,36–38]. Depletion of PABPC1 in mammalian cells had minimal effects on transcriptome-wide translation efficiency [39,40], suggesting that the stimulatory effect of PABPC on translation initiation may be restricted to circumstances where translation initiation efficiency is rate-limited.

In addition to its interactions with initiation factors, PABPC also interacts with the release factor eRF3 via its PAM2 motif in metazoans and its P-C domains in yeast [2,6,41–44]. Translation termination involves stop codon recognition in the ribosomal A site and nascent peptide release by eRF1, whose hydrolysis function and conformational change are stimulated by eRF3’s GTPase activity [45,46]. PABPC is thought to enhance termination efficiency by promoting the recruitment of the eRF1-eRF3 release factor complex to the stop codon. The role of PABPC in termination is also inferred from: i) the observation that tethering PABPC1 or Pab1 downstream of premature termination codons (PTCs) antagonized nonsense-mediated mRNA decay (NMD), an mRNA decay pathway thought to be activated by the reduced termination efficiency of premature translation termination [47,48] and ii) the increased termination efficiency observed with proximity of a stop codon to the mRNA 3’ end [49,50]. Direct evidence for PABPC’s ability to enhance termination includes: i) addition of PABPC1 to an *in vitro* termination assay improved termination efficiency [51], ii) PABP-interacting protein PAIP1 and PAIP2 competed with eRF3 for free PABPC binding, reducing termination efficiency of PTCs *in vitro* [52], and iii) deletion of *PAB1* or Pab1’s P-C domains *in vivo* increased stop codon readthrough efficiency of reporter PTCs in a proximity-dependent manner [50]. However, as with initiation, a full understanding of PABPC’s role during termination of endogenous mRNAs *in vivo* is still lacking.

PABPC’s involvement in many major stages of mRNA regulation and translation complicate attempts to define a specific role for PABPC *in vivo* by deleting, depleting, overexpressing, or mutating the protein and have led to conflicting models of PABPC’s function. Therefore, to specifically assess direct and indirect consequences of deleting PABPC, we generated and analyzed RNA-Seq, ribosome profiling [53], and mass spectrometry data from yeast cells lacking Pab1. As expected, mRNA and protein abundance changed substantially in *pab1Δ* cells. We found that deleting *PAB1* resulted in a translation termination defect that is not likely due to reduced eRF1 and eRF3 protein levels but rather appears attributable to loss of Pab1’s stimulatory function on termination. In addition, translation initiation defects and changes in relative translation efficiency in *pab1Δ* cells may be confounded by reduced initiation factor levels, especially eIF4G and eIF1. Further, an analysis of decapping activator substrates revealed that increased levels of certain mRNA subgroups may be partially caused by reduced levels of a specific decapping activator or components of Ccr4-Not deadenylation complex. Together, our results demonstrate direct and indirect consequences of *PAB1* deletion and illustrate the complexity of Pab1’s role in transcriptome-wide regulation of translation and mRNA decay.

## RESULTS

### Deletion of *PAB1* promotes transcriptome-wide accumulation of ribosomes downstream of normal stop codons

Deletion of *PAB1* has been shown to decrease translation termination efficiency and increase stop codon readthrough efficiency of reporter PTCs *in vivo* [50]. PTC readthrough occurs when a near-cognate tRNA outcompetes eRF1 in stop codon decoding, resulting in continued in-frame translation elongation and production of a C terminally extended polypeptide [54,55]. To determine whether decreased termination efficiency and increased readthrough efficiency could also be observed at normal termination codons (NTCs) of endogenous mRNAs when Pab1 is absent, we performed ribosome profiling and RNA-Seq analyses of yeast cells harboring a *PAB1* deletion. Because *PAB1* is an essential gene, the deletion was done in a *pbp1Δ* background, which suppresses *pab1Δ* lethality [56], and we used the *PAB1/pbp1Δ* strain as our wild-type *PAB1* control. Detailed descriptions of *pbp1Δ* cells have been described elsewhere [56–58]. Sequencing data obtained from three biological replicates of each strain were reproducible, as evidenced by high Pearson’s correlation coefficients between replicates (S1A Fig). In all strains, P-sites of ribosome profiling reads mostly map to the coding region of mRNAs, correspond to one major reading frame (frame 0), and show the expected 3-nucleotide periodicity indicative of translating ribosomes (Fig 1A-C, “CDS” region). Notably, the relative amount of ribosome footprints found in mRNA 3’-UTR regions is increased in *pab1Δpbp1Δ* cells compared to *pbp1Δ* or WT cells (Fig. 1A inset and B), demonstrating that cells lacking Pab1 manifest an apparent termination defect. This defect occurred not only at canonical stop codons, as evidenced by an increase of footprints in the “extension” region (the 3’-UTR region from the canonical stop codon to the next in-frame stop codon), but also at the first in-frame stop codons downstream of the canonical stop codon, as manifested by an increase of footprints in the distal 3’-UTR region (Fig 1B).

**Fig 1.**
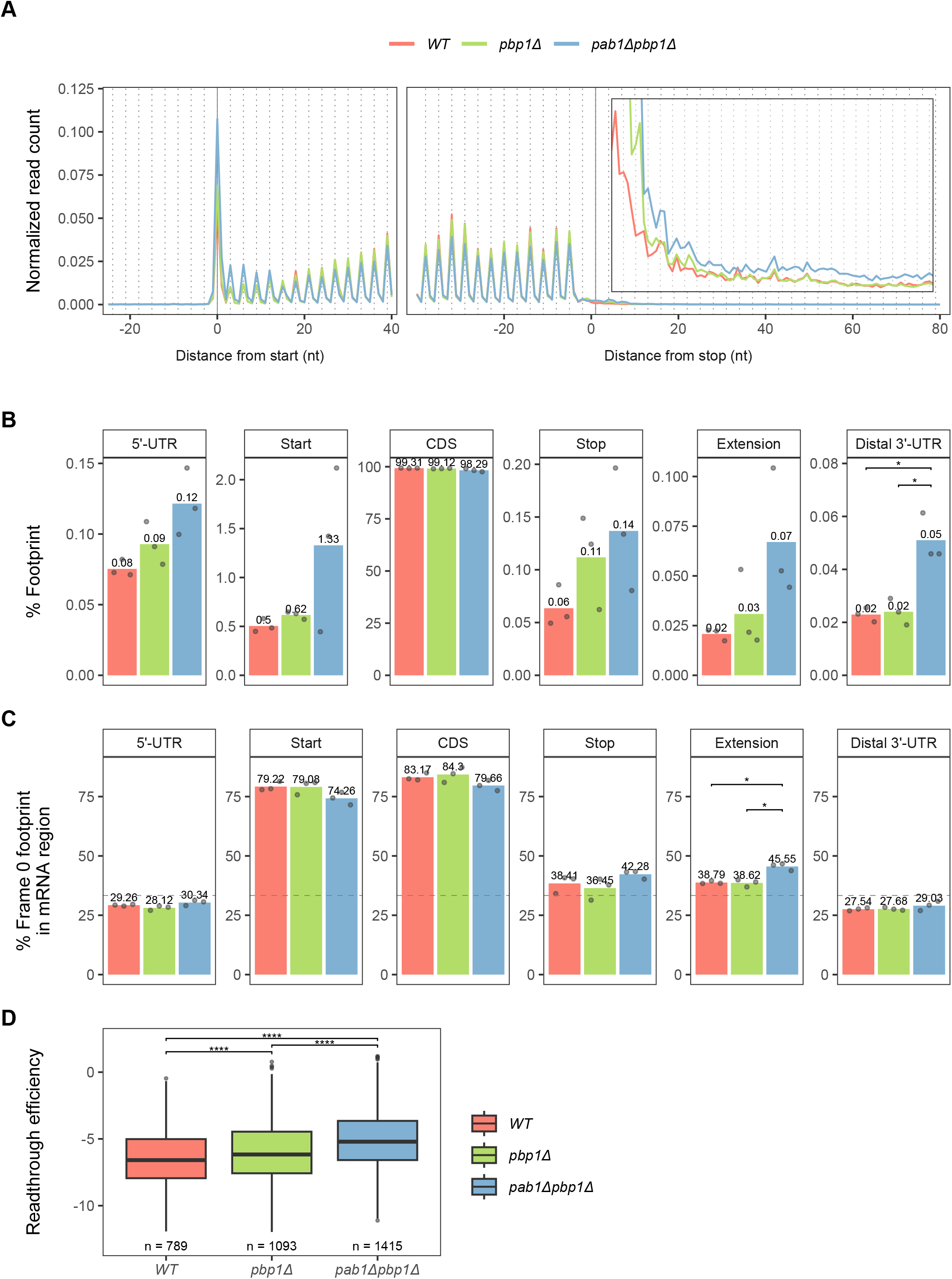
*PAB1* gene deletion leads to ribosome accumulation at mRNA start codons and in UTR regions. **A.** Ribosome footprints (replicate libraries were pooled) were counted by their P-site positions in the indicated nucleotide window around the canonical start and stop codons of annotated ORFs. Raw footprint counts were normalized by the total footprint count in the windows. Inset: Magnified view of the 3’-UTR region (Distance from stop > 0 nt). **B-C.** Percentage of footprints in sequencing library belonging in different mRNA regions (**B**) and percentage of frame 0 footprints in each mRNA region, where grey dashed line indicates a theoretical 33% at which all 3 reading frames are equally represented (**C**). “Start” region includes the canonical AUG and 3 flanking nucleotides on each side. “Stop” region includes the canonical stop codon and 3 flanking nucleotides on each side. “Extension” is the region following the “Stop” until (but not including) the first in-frame stop codon in the 3’-UTR. “Distal 3’-UTR” is the 3’-UTR region following “Extension.” Percentages from individual replicate libraries (grey points) were averaged (bar plot and reported value above it). Unpaired Student’s t-test with Benjamini-Hochberg method for multiple-testing correction was used to compare values between pairwise strains. **D.** Readthrough efficiency distribution in each strain (see Methods for calculation). Footprints from replicate libraries were pooled. Two-sided Wilcoxon’s rank sum test with Benjamini-Hochberg method for multiple-testing correction was used to compare values between pairwise strains. For B-D, only significant comparisons were reported as the following: (*) p < 0.05, (**) p < 0.01, (***) p < 0.001, (****) p < 0.0001. For all panels, only reads belonging to genes with UTR annotations and minimally overlapping sequences (less than 18 bp overlap with another gene on the same strand) were included in the analyses (2,693 genes).

The presence of ribosomes in the 3’-UTR can arise from stop codon readthrough, ribosome frameshifting, or reinitiation. To determine the primary driver of increased ribosome footprints in the 3’-UTR of mRNAs in *pab1Δpbp1Δ* cells, analyses of reading frame proportions in different mRNA regions were carried out. Stop codon readthrough would yield footprints predominantly in reading frame 0 in the extension region, while frameshifting or reinitiation events would not show this bias. Indeed, the proportion of ribosome footprints in reading frame 0 increases in the extension region in *pab1Δpbp1Δ* cells compared to *pbp1Δ* or WT cells (Fig 1C) and this increase is comparable to that observed in ribosome profiling data of cells depleted of functional eRF1 [59,60]. These results demonstrate that a notable portion of 3’-UTR footprints in *pab1Δpbp1Δ* cells arises from stop codon readthrough, thus suggesting that translation termination is less efficient in the absence of Pab1.

To verify that the increase in stop codon readthrough is transcriptome-wide, i.e., that the results of Figures 1A-C were not derived from a limited number of mRNAs, we calculated readthrough efficiency for each mRNA by dividing the density of frame 0 footprints in the extension by that in the CDS region. The number of mRNAs with detectable readthrough is increased in *pab1Δpbp1Δ* cells, almost double that observed in WT, and overall readthrough efficiency in *pab1Δpbp1Δ* cells is significantly higher than in the other two strains (Fig 1D).

Curiously, the *pbp1Δ* strain also shows higher overall readthrough efficiency than the WT strain (Fig 1D), raising the question of whether Pbp1 plays a role in termination and contributes to increased readthrough in the *pab1Δpbp1Δ* strain. However, this observation is most likely not due to stop codon readthrough. Footprints in the 3’-UTR region in the *pbp1Δ* strain are only higher than in WT in the few codons immediately following the canonical stop codon, while in the *pab1Δpbp1Δ* strain, the increase in 3’ UTR footprints extend substantially beyond that (Fig 1A, inset). More importantly, the proportion of frame 0 footprints in the extension region in *pbp1Δ* cells does not differ from that in WT cells (Fig 1C). Although we limited our readthrough efficiency calculation to only frame 0 footprints to ensure as accurate a calculation as possible, we still cannot completely exclude other events, such as reinitation, that happened to also produce ribosome footprints in frame 0. Thus, despite the fact that deletion of *PBP1* did not result in proportionally higher stop codon readthrough events compared to WT, a slight increase in footprints immediately following the stop codon leads to an apparent increase in the calculated readthrough efficiency values in the *pbp1Δ* strain.

In short, *PAB1* deletion resulted in a translation termination defect, manifested as increased footprints in mRNA 3’-UTR regions. The preference for frame 0 footprints in the extension region and the higher median readthrough efficiency values calculated for individual mRNAs observed in *pab1Δpbp1Δ* cells compared to the two controls indicate that stop codon readthrough increases transcriptome-wide in the absence of Pab1.

### Depletion of release factors is not likely to account for the effect of *PAB1* deletion on termination

Because PABPC plays major roles in the regulation of mRNA decay and translation, the termination defect and increased readthrough observed in *pab1Δpbp1Δ* cells can be due to: i) loss of Pab1’s direct function in termination via its interaction with eRF3 [44,51] or ii) changes in the stability of release factor mRNAs or changes in their translation that result in depletion of the respective proteins, which in turn affect global termination efficiency. To test for the latter scenario, we analyzed mRNA and protein abundance changes between strains using RNA-Seq and mass spectrometry data. As expected of Pab1’s essential and extensive role in mRNA stability regulation, steady-state mRNA and protein levels changed drastically when Pab1 was absent (Figs 2A, 2B, S2A, and S2B) while the absence of Pbp1 had minimal effects (S2A and S2B Fig). For proteins that were detectable by mass spectrometry, changes in their mRNA and protein abundance are quite consistent with each other, with Spearman’s correlation of 0.62-0.74 (Figs 2C and S2C). To study the effects of *PAB1* deletion, we focused further analyses on abundance changes in *pab1Δpbp1Δ* cells relative to *pbp1Δ* cells.

**Fig 2.**
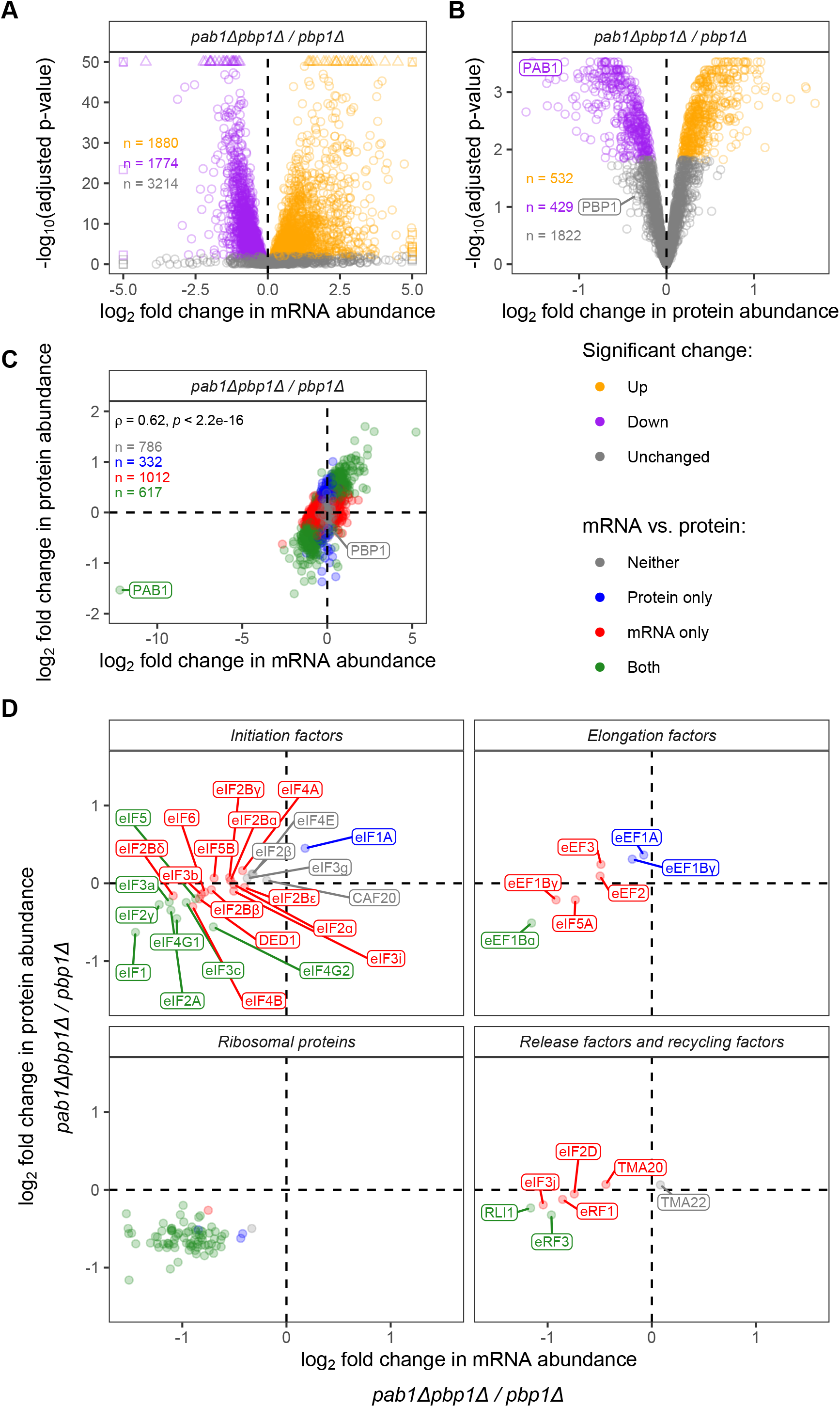
*PAB1* gene deletion has significant effects on the transcriptome and the proteome. **A.** Volcano plot of changes in transcriptome (RNA-Seq data) between *pab1Δpbp1Δ* and *pbp1Δ* strains. Orange, purple, and grey dots represent mRNAs with higher abundance (positive log_2_ fold change, adjusted p-value < 0.01), lower abundance (negative log_2_ fold change, adjusted p-value < 0.01), and no change (adjusted p-value ≥ 0.01), respectively, in the *pab1Δpbp1Δ* strain. **B.** Volcano plot of changes in proteome (mass spectrometry data) between *pab1Δpbp1Δ* and *pbp1Δ* strains. Orange, purple, and grey dots represent proteins with higher abundance (positive log_2_ fold change, adjusted p-value < 0.015), lower abundance (negative log_2_ fold change, adjusted p-value < 0.015), and no change (adjusted p-value ≥ 0.015), respectively, in the *pab1Δpbp1Δ* strain. **C.** Comparison of log_2_ fold change in transcriptome and proteome, with Spearman’s correlation coefficient. Grey, genes whose mRNA and protein abundance remained unchanged. Blue, genes whose protein but not mRNA abundance changed significantly. Red, genes whose mRNA but not protein abundance changed significantly. Green, genes whose mRNA and protein abundance both changed significantly. **D.** As in C, with the focus on translation-related genes. The ribosomal proteins depicted here account for 94% of all ribosomal proteins that make up the 40S and 60S subunits.

Among the detectable proteins related to mRNA translation, the *pab1Δ* mutation had the strongest effect on the levels of ribosomal proteins (Fig 2D, bottom left) and multiple initiation factors (Fig 2D, top left), but only appeared to have small effects on elongation factors (Fig 2D, top right). This result is corroborated by gene ontology analysis, which identified translation and ribosome biogenesis as two of the many pathways enriched in the down-regulated group of proteins (S1 Table and S1 file).

For the two release factors, eRF1 (Sup45) showed a slight reduction in its mRNA level but not its protein level, while eRF3 (Sup35) showed significant reduction in both mRNA and protein levels in the absence of Pab1 (Fig 2D, bottom right), where the eRF3 protein level was 80% of that in *pbp1Δ* cells. However, the reduction in release factor levels may not necessarily result in reduced termination efficiency. If overall translation is also reduced, the normal stoichiometry of supply and demand for release factors may still be maintained or supply may even exceed demand. The latter conclusion follows from observations that PABPC depletion substantially reduces overall protein synthesis such that almost all heavy polysomes are lost [40,56] and protein synthesis is limited by the amount of free ribosomes [30,61–63]. Hence, since termination can only occur after initiation and elongation, the larger reductions in ribosomal proteins and initiation factors (the most reduced ribosomal protein and initiation factor are respectively reduced to 45% and 64% of their normal levels) would most likely be limiting and release factors would thus be expected to still be in excess.

We also considered whether increased ribosome footprints in the 3’-UTR arise from a reduced level of Rli1, a ribosome recycling factor. However, since recycling can only occur after termination, Rli1 may still be in excess stoichiometrically in the context of overall reduced translation in *pab1Δpbp1Δ* cells. More importantly, *pab1Δpbp1Δ* cells show a preference for frame 0 in the extension region (Fig 1C), consistent with stop codon readthrough, while Rli1 depletion cells resulted in reinitiation in the 3’-UTR in all three reading frames [64].

Together, previous studies and our results here suggest that a significant reduction in translation is reducing the demand for release factors and recycling factors, making them likely to still be in excess. Therefore, the observed decreased termination efficiency or increased stop codon readthrough in *pab1Δpbp1Δ* cells is likely due to loss of Pab1’s stimulatory function on termination.

### 3’-UTR length is no longer predictive of readthrough efficiency when *PAB1* is deleted

Pab1 has been shown to affect NMD-sensitivity and readthrough efficiency of PTC containing mRNAs in a manner dependent on PTC proximity to mRNA-associated Pab1 [47,50,65]. As would be expected from this relationship, readthrough of PTCs in reporter mRNAs increased in response to 3’-UTR lengthening, but this trend was lost when *PAB1* was deleted [50]. Recently, we investigated the *cis*-regulatory elements of transcriptome wide stop codon readthrough using a random forest machine learning approach and found that 3’-UTR length was an important predictor of readthrough, where mRNAs with short 3’-UTRs had lower readthrough than those with long 3’-UTRs when eRF1’s functionality was compromised, but the trend was the opposite in WT cells [59]. These data led us to further assess the involvement of Pab1 and its proximity to the stop codon as a predictor of readthrough efficiency. If Pab1 is involved, we expected that the relationship between readthrough efficiency and 3’-UTR length would disappear or weaken when *PAB1* is deleted. Thus, we applied the same random forest approaches to identify mRNA features that influence the prediction of readthrough efficiencies in WT, *pbp1Δ*, *pab1Δpbp1Δ* strains (Figs 3A and S3A). As expected, the negative control features (NC) have no influence on readthrough efficiency prediction in any strain, while the identity of the stop codon is an important predictor of readthrough in all strains (Fig 3A). The length of the 3’-UTR is an important predictor of readthrough efficiency in WT and *pbp1Δ* cells but is no longer important in *pab1Δpbp1Δ* cells (Fig 3A). The relationship between 3’-UTR length and readthrough efficiency is slightly negatively correlated in WT and *pbp1Δ* strains, but it is weakened in the *pab1Δpbp1Δ* strain (Fig 3B, “all”). The fact that the correlation is not completely eliminated even in the absence of Pab1 could be due to technical limitations of readthrough efficiency calculation, where frame 0 non readthrough 3’-UTR footprints were inevitably included. Nevertheless, when other mRNA features were included in the analysis and controlled for (i.e., random forest regression model), this weak correlation became insignificant in predicting readthrough efficiency in *pab1Δpbp1Δ* cells.

**Fig 3.**
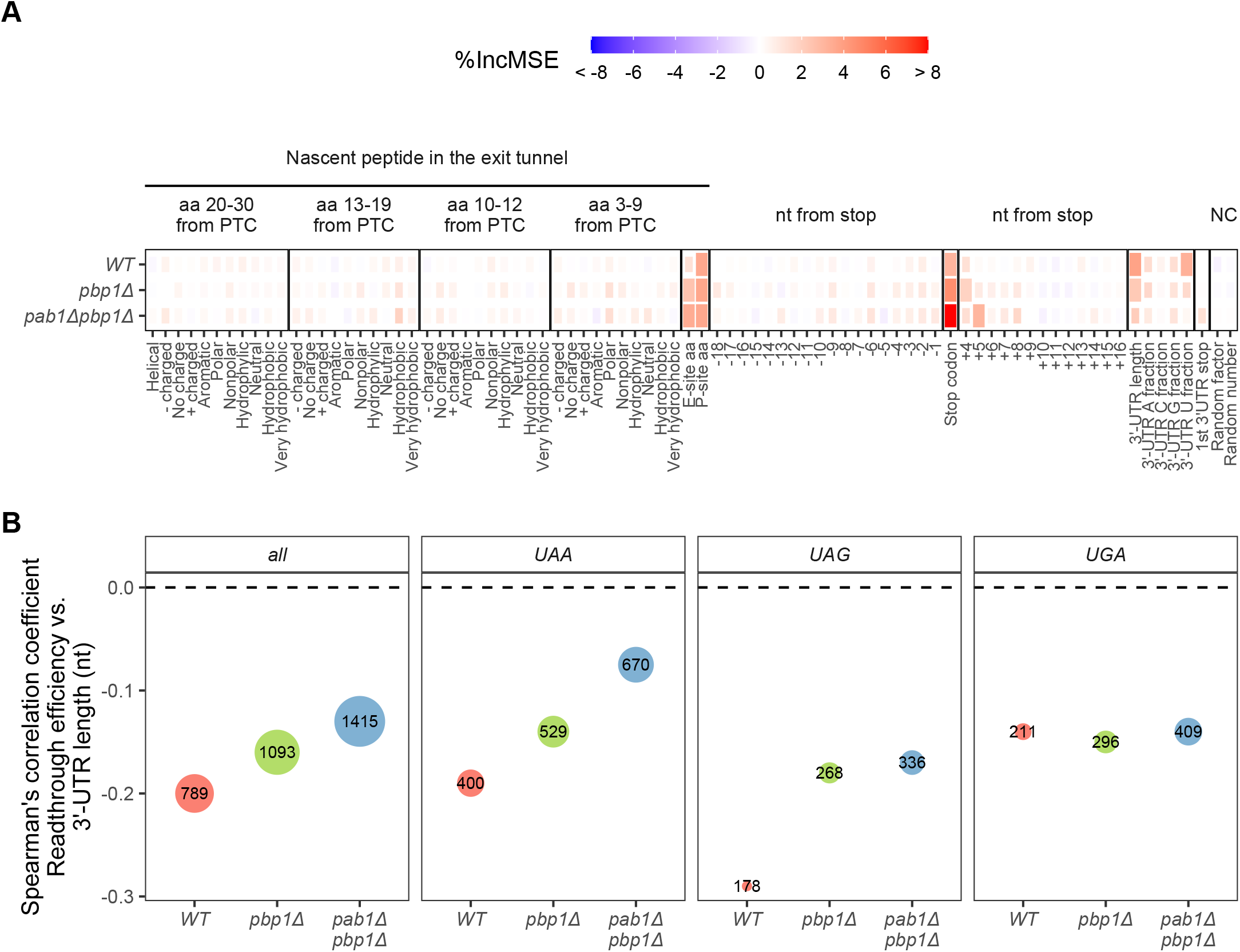
Pab1 regulates readthrough efficiency in a distance-dependent manner. **A.** Average feature importance scores (percent increase in mean squared error (%IncMSE)) extracted from 25 random forest models (5-fold cross-validation, repeated 5 times) trained for each strain to predict readthrough efficiency. Higher feature importance score (red) means the prediction error is high when that feature is permuted. As negative controls (NC), each mRNA was assigned arbitrary continuous and discrete values (Random number and Random factor). Features with significant importance (empirical p-value < 0.05 in at least 15 out of 25 models) are represented as bigger tiles. **B.** Spearman’s correlation coefficient of the relationship between readthrough efficiency and 3’-UTR length using all available data (“all”) or split data by stop codon usage. Number and dot size reflect the number of mRNAs in each correlation coefficient.

To further see how 3’-UTR length synergistically regulates readthrough with the strongest feature, stop codon identity, we grouped mRNAs by their stop codon identities and then performed the correlation analysis (Fig 3B). We found that the correlation between readthrough efficiency and 3’-UTR length is closer to zero in the absence of Pab1 than those observed in the other two strains for mRNAs with UAA as the stop codon, which happens to be the most common stop codon in the yeast transcriptome, but this trend isn’t observed for UAG and UGA (Fig 3B). This result suggests that mRNAs with UAG and UGA, although allowing higher readthrough (Fig S3C), are less sensitive to the proximity of Pab1 to the stop codon in this readthrough measurement, possibly because termination is slower and rate-limiting, unlike UAA where termination is faster. Since stop codon identity is a more important feature affecting readthrough efficiency, the effect caused by loss of Pab1 is somewhat masked.

Overall, we find that the proximity of Pab1 to the stop codon, as measured by 3’ UTR length, plays a role in readthrough efficiency prediction in combination with other known mRNA features, and the predictive ability of 3’-UTR length is lost in *pab1Δpbp1Δ* cells. Because 3’-UTR length was still predictive of readthrough efficiency in cells depleted of functional eRF1 [59], the fact that it no longer predicts readthrough of mRNAs in *pab1Δpbp1Δ* cells further supports the notion that readthrough occurring in *pab1Δpbp1Δ* cells is likely due to loss of Pab1’s direct function in termination rather than reduction in functional release factor levels.

### Deletion of *PAB1* leads to translation initiation defects but has minimal effects on translation efficiency

Pab1 is thought to promote translation initiation by aiding the association of the 40S ribosomal subunit and the eIF4F complex with the mRNA 5’ cap through a direct interaction with eIF4G [2,26,29,30]. Translation initiation defects may thus be expected when *PAB1* is deleted and, consistent with this hypothesis, we observed increased accumulation of ribosomes at the canonical AUG start codon and in the 5’-UTR region in the *pab1Δpbp1Δ* strain (Fig 1A and 1B).

To determine whether deletion of *PAB1* affected initiation rates of all mRNAs equally, we assessed the change in relative translation efficiency (TE) of each mRNA between *pab1Δpbp1Δ* and *pbp1Δ* strains (Fig 4A). Although ∼500 mRNAs show substantive increases or decreases in TE, most mRNAs (>90% of the transcriptome) do not show significant changes in relative TE (Fig 4A, grey). These results indicate that the absence of Pab1 affects the initiation process of most mRNAs to the same extent, such that the number of ribosomes recovered for a particular mRNA ORF remains proportional to the mRNA level. This observation is consistent with previous reports of human cells depleted of PABPC [39,40]. However, we cannot rule out an indirect effect of *PAB1* deletion on translation initiation factor levels. As shown in the analysis of transcriptome and proteome changes in response to *PAB1* deletion, some initiation factors were reduced at both mRNA and protein levels (Fig 2D). These disproportionate initiation factor levels could potentially cause a global reduction in translation and the appearance of unchanged relative mRNA TE. Notably, among the most reduced initiation factors are the two paralogs of eIF4G (eIF4G1/Tif4631 and eIF4G2/Tif4632), a binding partner of Pab1.

**Fig 4.**
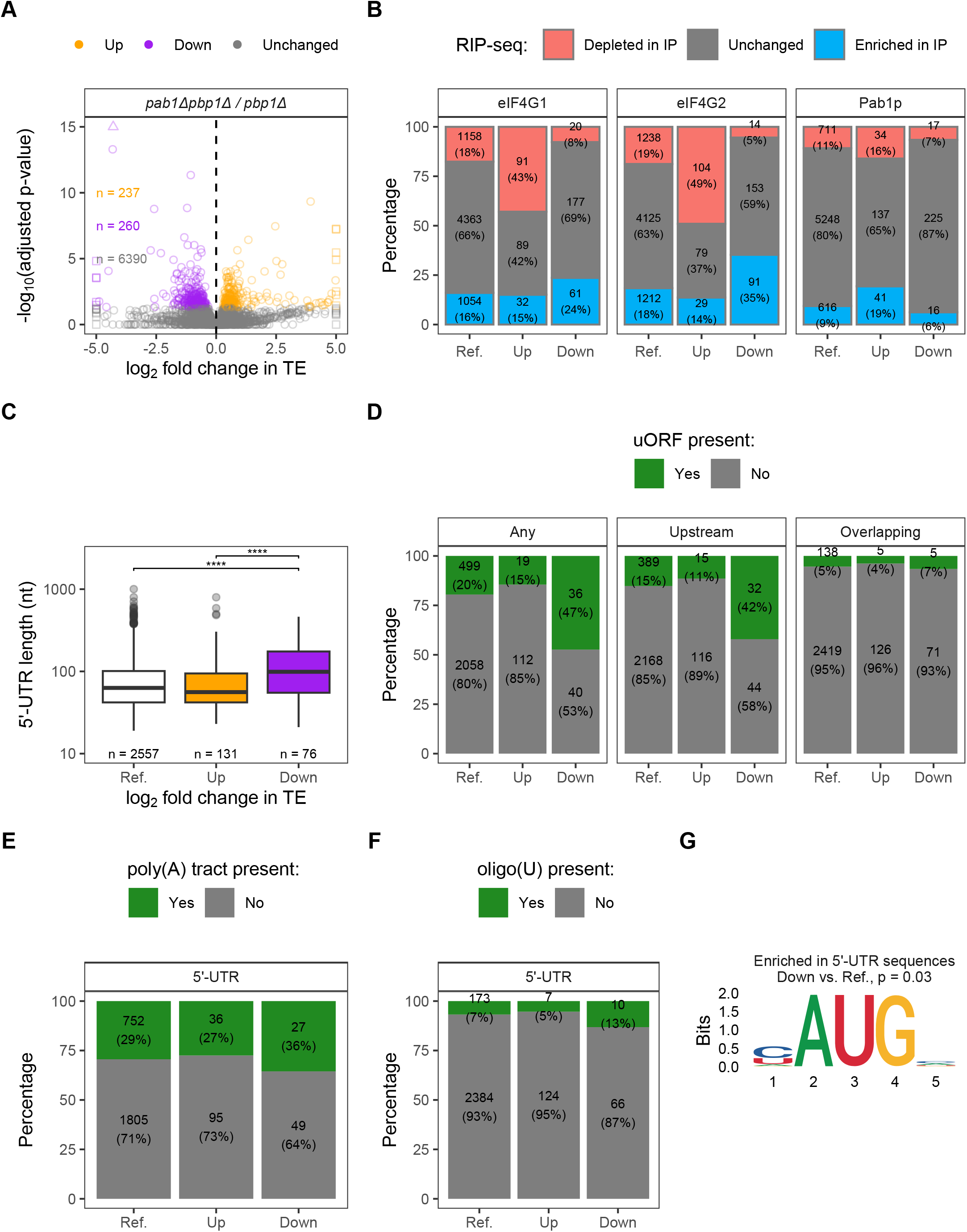
Translation initiation defects in response to *PAB1* deletion are indirect. **A.** Volcano plot of changes in translation efficiency (TE) between *pab1Δpbp1Δ* and *pbp1Δ* strains. Orange, purple, and grey dots represent mRNAs with increased (positive log_2_ fold change, adjusted p-value < 0.05), decreased (negative log_2_ fold change, adjusted p-value < 0.05), and unchanged TE (adjusted p-value ≥ 0.05), respectively, in the *pab1Δpbp1Δ* strain. **B.** Proportion and number of mRNAs from groups in A that were enriched (positive log_2_ fold change, FDR < 0.05), depleted (negative log_2_ fold change, FDR < 0.05), or unchanged (FDR ≥ 0.05) in RIP-seq data [36]. Pairwise ξ^2^ test with Benjamini-Hochberg method for multiple-testing correction was used to compare between Reference, Up, and Down groups. p < 0.05 in all pairwise comparisons (exact values provided in S2 Table) **C.** Distribution of 5’-UTR length in each mRNA group from A. Two-sided Wilcoxon’s rank sum test with Benjamini-Hochberg method for multiple-testing correction was used to compare values between pairwise groups. Only significant comparisons were reported as the following: (*) p < 0.05, (**) p < 0.01, (***) p < 0.001, (****) p < 0.0001 **D-F.** Proportion and number of mRNAs from groups in A with (“Yes”) or without (“No”) uORF (**D**), poly(A) tract (**E**), or oligo(U) (**F**) in the 5’-UTR. Pairwise Fisher’s exact test with Benjamini-Hochberg method for multiple-testing correction was used to compare between Reference, Up, and Down groups. p-values are provided in S2 Table. **G.** Motif enriched in 5’-UTR sequences of mRNAs in the Down group relative to Reference, identified by STREME. For C-G, analyses were limited to mRNAs with existing UTR annotations. Reference (Ref.) group includes all mRNAs regardless of TE changes (Up + Down + Unchanged) to recapitulate the general distribution of measured values in the transcriptome.

### *PAB1* deletion impacts differential translation initiation indirectly through the reduction of initiation factor levels

To further investigate how deletion of *PAB1* impacts translation initiation, we focused on 470 completely spliced mRNAs that showed significant increases or decreases in TE in response to *PAB1* deletion and compared them with two published data sets. First, we compared the identities of mRNAs that showed increased (“Up”) or decreased (“Down”) TE in our dataset to microarray data of polysome-associated mRNAs obtained from cells depleted for eIF4G [66]. For the mRNAs in the Up group, only 3 (1%) overlap between the two datasets (212 from our dataset and 99 from their dataset). For the mRNAs in the Down group, none overlap between the two datasets (258 from our dataset and 94 from their dataset). This lack of overlapping results could mean that these changes in TE are truly not through reduction of eIF4G or could be due to differences in experimental approaches, e.g., detection by microarray vs. RNA-Seq and perhaps the slight reduction in eIF4G does not give the same effect as complete depletion of the protein. For the second analysis, we utilized RIP-seq data of mRNAs associated with immunoprecipitated (IP’d) TAP-tagged eIF4G and Pab1 [36]. If our data was indirectly influenced by the reduction of eIF4G level, we would see that mRNAs enriched in IP of eIF4G and therefore highly dependent on eIF4G for translation initiation would be most sensitive to eIF4G reduction – they would have decreased TE and therefore be found in our Down group. Indeed, we found that mRNAs that were enriched in IPs of eIF4G1 or eIF4G2 are over-represented in the Down group compared to the general distribution in the transcriptome (Reference “Ref.” group) (Fig 4B, red, and S2 Table). Moreover, mRNAs that were depleted in IPs of eIF4G1 or eIF4G2, that may rely less on cap dependent initiation, are over-represented in the Up group and under-represented in the Down group (Fig 4B, blue, and S2 Table). Notably, this trend is stronger than what is observed in Pab1 IP (Fig 4B, and S2 Table). Overall, these results indicate that changes in TE when *PAB1* is deleted are at least partially mediated by a reduction in eIF4G levels.

Despite the disparate effect of eIF4G on different mRNA subgroups, eIF4G still does not explain the entirety of the data. For example, although the mRNAs enriched in eIF4G2 IP are proportionally over-represented in the Down group compared to Reference (35% vs. 18%), 65% of the mRNAs in the Down group are those not highly dependent on eIF4G (Fig 4B). Thus, we further characterized these mRNAs by exploring their 5’-UTR features.

First, we observed that mRNAs in the Down group tend to have longer 5’-UTRs than those in the Up or Reference groups (Fig 4C). A longer 5’-UTR increases the chance for motifs or upstream ORFs (uORFs) that could interfere with initiation complex assembly or the scanning mechanism. The presence of uORFs is generally thought to suppress translation initiation of the main ORF [67]. We found that the proportion of mRNAs with at least one uORF is higher in the Down group than in other groups (Fig 4D, left panel, and S2 Table). We further asked if the results were due to uORFs that are completely upstream or uORFs that are overlapping with the main ORF by conducting the same analysis for the two types of uORFs separately. The presence of overlapping uORFs is quite rare in our data set and the proportions are not different between groups, but the results for upstream uORFs mimicked those of all uORFs analyzed together (Fig 4D and S2 Table). The sensitivity of uORF-containing mRNAs to *PAB1* deletion could be related to the reduction in eIF1 (Sui1) level in the *pab1Δpbp1Δ* strain (Fig 2D). eIF1 has a role in start codon recognition, discriminating against suboptimal start sites in favor of the optimal one [68–71]. Therefore, it is not surprising that mRNAs with decreased TE tend to have uORFs (Fig 4D). Consistent with this notion, ribosome footprints in the 5’-UTR region are increased in the *pab1Δpbp1Δ* strain relative to the other two strains (Fig 1B).

Next, we considered motif-dependent regulation of translation initiation. First, recruitment of Pab1 to poly(A) tracts in the 5’-UTR was reported to induce internal cap independent initiation in yeast [72]. Therefore, mRNAs with this motif would be expected to have decreased TE in the absence of Pab1 and thus be found more often in our Down group. However, we did not find differences in the proportions of mRNAs containing poly(A) tracts in the 5’-UTR between groups (Fig 4E and S2 Table). Second, oligo(U) longer than 7 nt in the 5’-UTR was identified as eIF4G1’s preferential binding motif and can promote initiation [73,74]. However, we did not find differences in the proportions of mRNAs containing oligo(U) in the 5’-UTR between groups (Fig 4F and S2 Table). In a parallel approach, we used the motif discovery tool STREME [75] to identify motif(s) enriched in 5’-UTR sequences of either Up and Down group relative to the Reference. Neither poly(A) tract nor oligo(U) is enriched in either group compared to sequences from the Reference group, consistent with our direct analyses (Fig 4E and 4F), but the AUG motif is enriched in the Down group (Fig 4G), consistent with our uORF analysis (Fig 4D). No other novel motif was identified.

We also asked whether nucleotide context around the start codon and near the 5’ cap influence changes in TE in response to *PAB1* deletion by comparing proportions of nucleotides at each position from each group relative to Reference (S4 Fig). The optimal context for translation in yeast has been determined as AA(A/G)A<u>AUG</u>UCU, with position -3 (3^rd^ nucleotide upstream of AUG, which is considered as positions +1 +2 +3) being the most important and conserved [67,76,77]. We did not find mRNAs in the Up or Down group to have biases in nucleotide usage at position -3 compared to Reference or each other (S4A Fig). However, other positions in this window that show significant differences relative to the Reference are consistent with the consensus, namely, A/C are enriched in the Up group at position -4 (S4A Fig) and U is depleted in the Down group at position +4 (S4B Fig). Beyond the immediate AUG context, G is enriched and U is depleted in the Down group at position +18 (S4B Fig). From the 5’ cap, U is enriched and A is depleted in the Down group at position +6 (S3C Fig).

Overall, our results show that studying Pab1’s role on global translation initiation through deleting or depleting Pab1 can be confounded by the reduction in initiation factor levels, especially eIF4G and eIF1.

### Properties of the mRNAs with differential TE in response to *PAB1* deletion support the notion that mRNA 5’ and 3’ ends communicate

Changes in an mRNA’s poly(A) tail length can change the extent of its commitment to translation initiation, observations which led to the model of 5’-3’ communication of mRNA ends in translation [18,19,23,24]. Pab1 is thought to play an important role in facilitating this closed-loop mRNA structure, bridging the interaction with both the poly(A) tail and eIF4G [27–29]. Shorter mRNAs are thought to form more stable structures than longer mRNAs [28,65], but the closed-loop structure may not apply to every mRNA as not all mRNPs contain the closed-loop components [36,38].

Although it is uncertain on which mRNAs the closed-loop occurs, the possibility and efficiency of 5’-3’ communication may still be explained through simple proximity of 5’ and 3’ ends [38]. Efficient 5’-3 communication allows efficient feedback of ribosomes recycled from termination to a new round of initiation, and this efficiency should be gene length-dependent, as diffusion of ribosomes between the ends would be expected to be more efficient for shorter mRNAs than longer mRNAs [62,63], even without Pab1 facilitating the closed-loop. This is especially relevant when the availability of ribosomes is limiting and ribosome recruitment becomes more dependent on recycled ribosomes than the limited free ribosomes [62,63], which may be the case in our data with the reduction in ribosomal protein levels in *pab1Δpbp1Δ* cells (Fig 2D). When comparing CDS and entire mRNA lengths between groups, we found that mRNAs in the Down group tend to be longer than both the Up group and the Reference while those in the Up group tend be shorter (Fig 5A and 5B). These results are consistent with the proximity-based 5’-3’ communication model.

**Fig 5.**
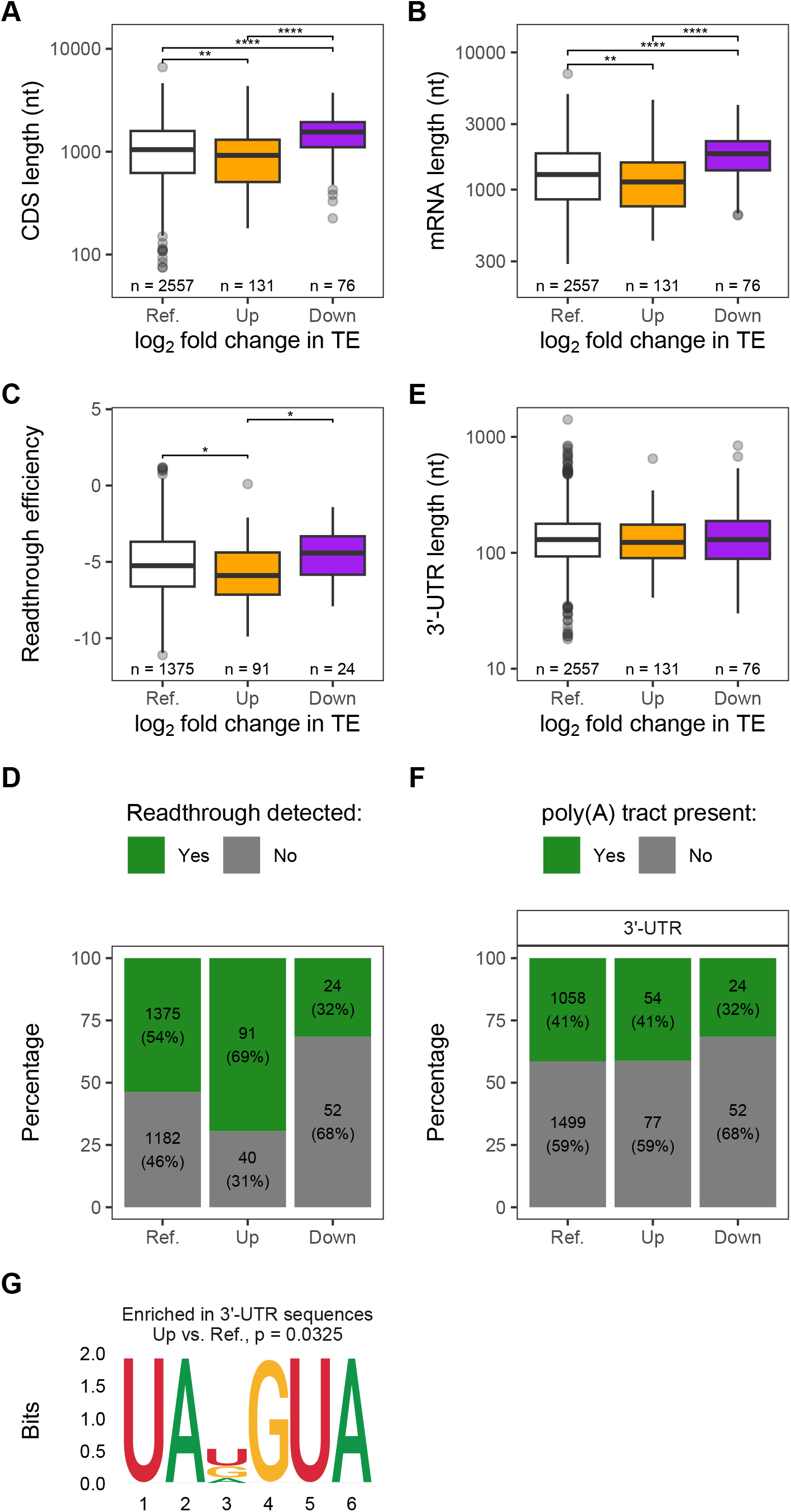
Evidence for communication of mRNA 5’-3’ ends in efficient initiation. **A-B.** Distribution of CDS length (**A**) or mRNA length (**B**) in each mRNA group from Fig. 4A. **C.** Distribution of readthrough efficiency for mRNAs with detectable readthrough in the *pab1Δpbp1Δ* strain in each group. **D.** Proportion and number of mRNAs with detectable (“Yes”) or not detectable (“No”) readthrough in the *pab1Δpbp1Δ* strain. **E.** Distribution of 3’-UTR length in each mRNA group. **F.** Proportion and number of mRNAs with (“Yes”) or without (“No”) poly(A) tracts in the 3’-UTR. For A, B, C, and E, two-sided Wilcoxon’s rank sum test with Benjamini-Hochberg method for multiple-testing correction was used to compare values between pairwise groups. Only significant comparisons were reported as the following: (*) p < 0.05, (**) p < 0.01, (***) p < 0.001, (****) p < 0.0001. For D and F, Pairwise Fisher’s exact test with Benjamini-Hochberg method for multiple-testing correction was used to compare between groups. p-values are provided in Supplemental Table S2. **G.** Motif enriched in the 3’-UTR sequences of mRNAs in the Up group relative to Reference, identified by STREME. For all panels, analyses were limited to mRNAs with existing UTR annotations. Reference (Ref.) group includes all mRNAs regardless of TE changes (Up + Down + Unchanged) to recapitulate the general distribution of measured values in the transcriptome.

Consistent with the notion that efficient recycling of ribosomes at the 3’ end promotes efficient translation initiation at the 5’ end, we found that mRNAs with increased TE have lower readthrough efficiency (i.e., more efficient termination) while those with decreased TE have higher readthrough efficiency (Fig 5C). We limited our analysis to mRNAs with detectable readthrough due to the cyclic nature of translation and the detection limit of readthrough ribosomes. Since the amount of readthrough ribosomes depends on the amount of translation of the CDS, for mRNAs with the same readthrough *efficiency*, readthrough may not be detectable for mRNAs with lower TE (so readthrough efficiency appears to be zero for them) but remain or become detectable for mRNAs with higher TE. Thus, the fact that the proportion of mRNAs with detectable readthrough in the Down group is lower than the Up group (Fig 5D) does not necessarily mean that readthrough efficiency is lower in the Down group. In sum, we found that mRNAs with increased TE had lower readthrough efficiency and those with decreased TE had higher readthrough efficiency (Fig 5C).

To see if Pab1’s function at termination influences the distinction between Up and Down TE groups in response to *PAB1* deletion, we compared 3’-UTR lengths among groups, but found no significant differences (Fig 5E). This result is perhaps not surprising, as 3’-UTR length is not the strongest feature in WT conditions and no longer matters in the *pab1Δpbp1Δ* strain (Fig 3A). We next asked whether a specific sequence motif is enriched in either Up or Down group. No differences between groups were detected in terms of the presence of poly(A) tracts in the 3’-UTR region (Fig 5F), which can serve as a binding site of Pab1 in addition to the poly(A) tail, and this result was confirmed by the lack of poly(A) motif in the motif discovery approach, STREME. However, STREME identified the UAKGUA motif enriched in the Up group relative to Reference (Fig 5G). The UA…UA sequence may indicate two consecutive strong stop codons, either UAA or UAG, where the second stop codon (although out-of-frame with the first) acts as a fail-safe stop codon in case of failed termination or recycling at the first stop codon. Moreover, the enrichment of G following the first stop codon is consistent with the observation that mRNAs with G at this position had the lowest readthrough efficiency (S3C Fig, “+4”). This motif being enriched in mRNAs in the Up group, along with the shorter mRNA length and lower readthrough efficiency of this group, is consistent with the hypothesis that more efficient ribosome recycling promotes efficient new rounds of translation initiation.

### Substrates of decapping activators Pat1/Lsm1 and Upf1/Upf2/Upf3 tend to be more increased than decreased in response to *PAB1* deletion

Among its many functions, PABPC also has important regulatory roles in mRNA decay [1–3]. Hence, we asked how the absence of Pab1 impacts the levels of mRNAs that are substrates of different decapping activators, namely Dhh1, Pat1/Lsm1, and the Upf factors of the NMD pathway. Dhh1, Pat1/Lsm1, and NMD substrates are defined respectively as mRNAs whose levels were increased in *dhh1Δ* cells, commonly increased in *pat1Δ* and *lsm1Δ* cells, and commonly increased in *upf1Δ, upf2Δ, and upf3Δ* cells, relative to WT [78,79]. Approximately half of the mRNAs in each group showed significant changes in abundance in *pab1Δpbp1Δ* relative to *pbp1Δ* cells. Of mRNAs that showed significant changes, those that are targets of either Pat1/Lsm1 or NMD or both tend to be increased rather than decreased (Fig 6A, top and middle panels, columns 3-8, compare Up (orange) to Down (purple)). On the other hand, mRNAs that are only substrates of Dhh1 are somewhat comparably increased and decreased (Fig 6A, top and middle panels, column 2, compare Up (orange) to Down (purple)).

**Fig 6.**
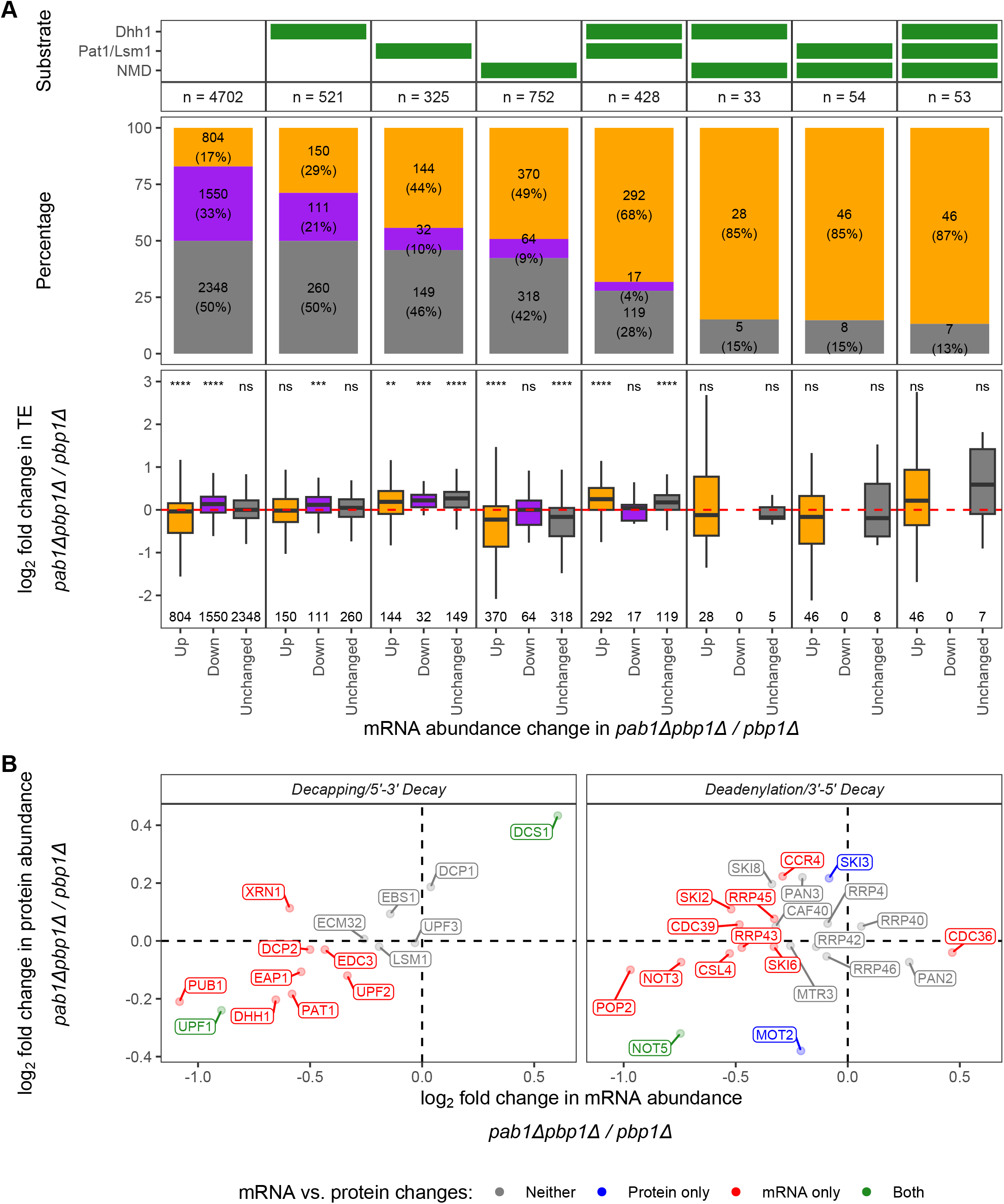
Changes in the abundance of decapping activator substrates by different consequences of *PAB1* deletion. **A.** mRNA abundance and translation efficiency changes between *pab1Δpbp1Δ* and *pbp1Δ* strains for Dhh1, Pat/Lsm1, and NMD substrates. ***(Top)*** Panel indicating substrate status (green) of panels below. A substrate is defined as an mRNA that is upregulated upon decapping deactivator gene deletion [9,79]. Dhh1: mRNAs upregulated in a *dhh1Δ* strain relative to WT [79]; Pat1/Lsm1: mRNAs commonly upregulated in *pat1Δ* and *lsm1Δ* strains relative to WT [79]; NMD: mRNAs commonly upregulated in *upf1Δ*, *upf2Δ*, and *upf3Δ* strains relative to WT [78]. ***(Middle)*** Proportions of mRNA abundance changes from Fig 2A: Up (orange), Down (purple), and Unchanged (grey), separated into columns by substrate status. ***(Bottom)*** Distribution of log_2_ fold change in TE between *pab1Δpbp1Δ* and *pbp1Δ* strains for mRNA groups from the middle panel. Two-sided Wilcoxon’s rank sum test with Benjamini-Hochberg method for multiple-testing correction was used to compare median log_2_ fold change in TE of zero (no change). Significant levels were reported as the following: (ns) not significant, (*) p < 0.05, (**) p < 0.01, (***) p < 0.001, (****) p < 0.0001. **B.** Comparison of log_2_ fold change in transcriptome and proteome as in Fig 2C, with the focus on factors related to mRNA decay.

To determine whether increases in the abundance of Pat1/Lsm1 and Dhh1 substrates, which follow the canonical deadenylation-dependent pathway, was due to loss of Pab1’s role in deadenylase recruitment or dysregulation of decay pathway components, we investigated changes in mRNA and protein levels of genes involved in decapping, deadenylation, and decay (Fig 6B). We found that while decapping enzyme (Dcp1 and Dcp2), Pat1, Lsm1, Dhh1, exonuclease Xrn1, and exosome components are not significantly enriched or depleted at the protein level, two proteins that are components of the Ccr4-Not deadenylase complex, Mot2 (also commonly known as Not4) and Not5, are significantly depleted to ∼77-80% of the amount in *pbp1Δ* cells (Fig 6B). Not4 and Not5 are thought to link slow translation elongation of non-optimal codons to deadenylation as well as deadenylation to decapping [3]. Specifically, Dhh1’s association with the ribosome requires Not5’s ribosome binding and Not4’s E3 ligase activity to ubiquitinate 40S ribosomal subunit protein eS7 [80]. Not5 has also been shown to bind Pat1 to promote decapping [81]. Thus, stabilization of Pat1/Lsm1 and some Dhh1 substrates may be partially attributable to dysregulated Ccr4-Not components, in addition to loss of Pab1’s role in deadenylase recruitment.

Because decapping of NMD substrates is usually a deadenylation-independent mechanism that is triggered by premature translation termination [9,82,83], stabilization of NMD substrates in *pab1Δpbp1Δ* cells is likely unrelated to Pab1’s role in protecting the poly(A) tail or deadenylase recruitment. Rather, it is likely due to decreases in translation, which leads to decreased frequency of premature termination. To test whether NMD substrates have a decreased initiation rate in the absence of Pab1, we compared log_2_ fold change in TE between *pab1Δpbp1Δ* and *pbp1Δ* strains for each group of mRNAs to the value of unchanged TE (log_2_ fold change = 0) (Fig 6A, bottom panel). Consistent with our hypothesis, we found that NMD substrates that are significantly stabilized or have unchanged mRNA levels in *pab1Δpbp1Δ* have relatively lower TE while the minority that are depleted have relatively unchanged TE (Fig 6A, bottom panel, column 4). However, reduced TE is probably not the only contribution to NMD substrate enrichment, as the protein level of the key NMD protein Upf1 is significantly reduced in *pab1Δpbp1Δ* cells to ∼85% of the level in *pbp1Δ* cells (Fig 6B).

## DISCUSSION

Numerous biochemical, structural, *in vitro*, and *in vivo* studies of specific mRNAs have identified pleiotropic roles for PABPC in cytoplasmic mRNA deadenylation, translation initiation, and translation termination [3]. However, because of PABPC’s multiple apparent functions, defining its transcriptome-wide roles has been difficult. High throughput approaches exploring transcriptome-wide effects of PABPC depletion have been carried out in mammalian cells [39,40], but none have been done in yeast. Consistent with observations in mammalian cells, we found that deletion of yeast *PAB1* resulted in major changes in the transcriptome (Fig 2A), but only minimal changes in relative translation efficiency (Fig 4A). Further, we showed that *pab1Δ* cells also drastically changed their proteome (Fig 2B) and provided the first evidence for a transcriptome-wide translation termination defect (Fig 1).

Our proteomics data have provided insights to the direct vs. indirect consequences of *PAB1* deletion and suggested a new layer of complexity to interpreting genome-wide gene expression alterations in PABPC-depleted cells. PABPC’s role in translation initiation has been elusive, partly because the extent of its activity is dependent on the stoichiometry between PABPC, poly(A) tracts, and basal translation levels [39], and these experimental conditions frequently vary between studies. Moreover, Pab1 appears to have preferential association with certain mRNAs [36]. Hence, it seems counterintuitive that depleting PABPC/deleting *PAB1* reduces translation overall, yet relative TE is unchanged for most mRNAs [39,40]. These observations are akin to the effects of depleting eIF4G in yeast [66]. Since *PAB1* deletion also reduced eIF4G mRNA and protein levels (Fig 2D), it is possible that effects attributed to the absence of Pab1 are due to dysregulation of eIF4G mRNA stability and the consequent reduction of eIF4G, which in turn reduced global translation initiation. This sequence of events is also likely for ribosomal proteins, as PABPC depletion has been shown to cause accelerated decay of mRNAs with short poly(A) tails [39], which usually are characteristics of highly expressed, highly translated mRNAs, including those encoding ribosomal proteins [3,84]. Even when we focused our analyses on mRNAs that did have significant changes in TE, where their drastic changes may be due to specific factors outside of the global regulators of translation initiation, their properties are still related to eIF4G-dependent pre-initiation complex recruitment, eIF1-mediated start codon recognition, and efficiency of ribosome recycling (Figs 4 and 5). Due to their intrinsic properties, these mRNAs with significant increase or decrease in TE are respectively more or less dependent on these processes than most mRNAs and are thus more sensitive to reduction in these factors in *pab1Δ* cells (Fig 2D). As a result, the direct role of Pab1 in initiation is masked by the changes in these core components of initiation, i.e., the translation initiation defects observed in *pab1Δ* cells are mostly indirect.

Similar reasoning might be applied to translation termination since there is a slight reduction in release factor levels in *pab1Δ* cells (Fig 2D). However, unlike initiation, it is less likely that this reduction masked Pab1’s direct role in termination or was the major reason for increased stop codon readthrough because: i) reduced initiation factors and ribosomal proteins are more limiting than release factors, skewing the usual stoichiometry of demand vs. supply for release factors towards the supply, ii) deletion of Pab1’s eRF3 interacting domain only (*pab1ΔC*), which affects termination but not mRNA decay [44], yields significant stop codon readthrough of reporter mRNAs [50], and iii) 3’-UTR length, which approximates the distance of Pab1 to the stop codon, lost its ability to predict readthrough efficiency in *pab1Δ* cells as opposed to WT (Fig 3A) or eRF1 mutant cells [59]. Nevertheless, it remains to be determined whether a 20% reduction in termination factors in a *PAB1* strain would yield the same termination defect as a *pab1Δ* mutation and whether *pab1ΔC* cells significantly change their release factor levels. Because near cognate tRNAs compete with release factors in stop codon decoding, changes in aminoacyl-tRNA levels, synthesis, and modifications should also be considered.

We showed that the sets of mRNA substrates of Pat1/Lsm1-mediated decay and NMD both manifest relative increases in abundance and thus likely to be partially stabilized in *pab1Δ* cells despite their different decay mechanisms, while substrates of Dhh1-mediated decay did not have a tendency to increase or decrease (Fig 6A). Stabilization of Pat1/Lsm1 substrates may be attributed to the loss of Pab1’s role in stimulating deadenylation of mRNAs usually subject to deadenylation-dependent decay. However, there is also a slight depletion of Ccr4-Not components, Not4 and Not5 (Fig 6B), and the extent of how much this depletion contributes to substrate stabilization is unknown. The consequences of Not4 and Not5 depletion may also apply to some Dhh1 substrates (i.e., those that are increased) but not all, possibly due to Dhh1’s involvement in multiple decapping complexes which can lead to alternative degradation pathways [9]. Some complexes may require Pab1’s role in deadenylase recruitment while for others, Dhh1’s communication with the Ccr4-Not complex may be enhanced by lower Pab1 level [85]. In contrast to Pat1/Lsm1 substrates, partial increases in the levels of NMD substrates most likely arises from the combination of decreased translation and decreased Upf1. The reduction in overall translation would result in fewer instances of termination and, together with the slight reduction in Upf1, would lower the likelihood that a nonsense-containing mRNA would be targeted by NMD.

Collectively, our observations of the indirect effects of *PAB1* deletion may help explain the discrepancies in previous studies of PABPC’s functions and our multi-omics data can be helpful resources for the design of future experiments involving genetic manipulation, depletion, or overexpression of PABPC.

## MATERIALS AND METHODS

### Yeast strain construction

Yeast strains used in this study, listed in S3 Table, are in the W303 background. Gene deletions were achieved by the PCR-based method [86] and high efficiency transformation [87] of fragments amplified by oligonucleotides listed in S4 Table, synthesized by Integrated DNA Technologies (IDT). Three biological replicates (isolates) of each mutant strain and three technical replicates of isogenic wild-type (WT) strain were used for all experiments.

The *pbp1Δ* strain was made by replacing the coding sequence of the *PBP1* gene in a WT strain (HFY114) with the *URA3* gene. The *URA3* cassette was obtained from plasmid HFSE1380 [79] by PCR using URA3_5F_v2 and URA3_3R_v2 primers. Homology arms flanking the *PBP1* coding sequence were amplified from genomic DNA by PCR using PBP1_5F, PBP1_5R_v2, PBP1_3F_v2, and PBP1_3R primers. DNA fragments consisting of the homology arms flanking the *URA3* cassette were constructed by PCR and transformed into competent WT yeast cells. Verification of successful gene replacement was confirmed by PCR followed by Sanger sequencing using primers listed in S4 Table.

Subsequent deletion of the *PAB1* gene from the *pbp1Δ* strain was carried out by replacing the coding sequence of *PAB1* with kanMX, resulting in a *pab1Δpbp1Δ* strain. The kanMX cassette was obtained from the pCAS plasmid (Addgene, #60847) by PCR using PAB1_KanMX_F and PAB1_KanMX_R primers. Homology arms flanking the *PAB1* coding sequence were amplified from genomic DNA by PCR using PAB1_5H_ext_F2, PAB1_5H_ext_R, PAB1_3H_ext_F, and PAB1_3H_ext_R primers. DNA fragments consisting of the homology arms flanking the kanMX cassette were constructed by PCR and transformed into competent *pbp1Δ* (KMY01) yeast cells. Verification of successful gene replacement was confirmed by PCR followed by DNA sequencing using primers listed in Supplemental Table S4 as well as by examining growth phenotype. Doubling time for these strains grown in YEPD at 30°C were 1.5 hours for WT and *pbp1Δ* strains and 3.5-4.5 hours for the *pab1Δpbp1Δ* strain.

### Cell growth and harvest

Cells were grown in 1 L of YEPD at 30°C with shaking. When the OD_600_ of the culture reached 0.6–0.8, cells were collected by rapid vacuum filtration, flash-frozen in liquid nitrogen in the presence of Footprinting Buffer (20 mM Tris-HCl pH 7.4, 150 mM NaCl, 5 mM MgCl_2_) plus 1% TritonX-100, 0.5 mM DTT, 1 mM phenylmethylsulfonyl fluoride (PMSF), and 1X protease inhibitors, and lysed in a Cryomill (Retsch) (5 Hz, 2 min; 10 Hz, 15 min). Cell lysates were clarified by ultracentrifugation in a Beckman Coulter Optima L-90K Ultracentrifuge at 18,000 rpm for 10 min at 4°C, using a 50Ti rotor. Centrifugation was repeated for the supernatant at 18,000 rpm for 15 min at 4°C. Lysates were stored at -80°C in aliquots.

### Ribosome profiling library preparation and sequencing

Ribosome profiling libraries were prepared as described previously [88]. Lysates were digested with RNase I (Invitrogen, #AM2294) for 1 hour at 25°C with shaking at 700 rpm, and the reaction was stopped using SUPERase-In RNase Inhibitor (Invitrogen, #AM2694). RNase I-treated lysates were then layered onto a 1 M sucrose cushion in Footprinting Buffer plus 0.5 mM DTT and centrifuged in a Beckman Optima TLX Ultracentrifuge at 60,000 rpm for 1 hour at 4°C using a TLA100.3 rotor to isolate 80S ribosomes. Ribosome-protected fragments (RPFs) were extracted from pelleted 80S ribosomes using a miRNeasy kit (QIAgen, #217004) following the manufacturer’s protocol for enriched recovery of small RNAs (<200 nt). RNAs larger than 200 nt, which include ribosomal RNAs (rRNAs) and other large RNAs, were discarded. Small RNAs were 3’ dephosphorylated and 5’ phosphorylated with T4 polynucleotide kinase (T4PNK, NEB, #M0201S), and purified with RNA Clean and Concentrator-5 (Zymo Research, #R1013) according to the manufacturer’s instructions that separately recover small and large RNA fractions. Large RNA fractions were discarded. Approximately 1 μg of small RNAs in 8.5 μl were incubated with 2 μl of QIAseq FastSelect –rRNA Yeast Kit (Qiagen, #334215) at 75°C, 2 min; 70°C, 2 min; 65°C, 2 min; 60°C, 2 min; 55°C, 2 min; 37°C, 2 min; 25°C, 2 min; 4°C, hold. This step hybridized any remaining rRNAs in the sample to the rRNA oligonucleotides, creating duplexes that would fail to ligate to sequencing adapters or fail to be reversed-transcribed into cDNA for sequencing library preparation. Sequencing libraries were prepared from the 10.5 μl reactions using the NEXTflex Small RNA-Seq Kit v3 (Perkin Elmer/Bioo Scientific, #NOVA-512-05) according to the manufacturer’s protocol, except for the RNA denaturation step (70°C, 2 min incubation) before 3’ 4N Adenylated Adapter ligation, which was skipped. Based on the manufacturer’s instructions for optimization, adapters were undiluted, PCR was performed for 15 cycles, and the library was purified using the manufacturer’s gel-free size-selection and cleanup protocol. Extra rounds of cleanup were performed if the amount of PCR primers was still high compared to the amount of library, as analyzed on a Fragment Analyzer. Three libraries were multiplexed according to NEXTflex’s recommended combinations of barcodes (index sequences) and sequenced (single-end, 75 cycles) in-house on Illumina NextSeq 500 or NextSeq 550 sequencers.

### RNA-Seq library preparation and sequencing

RNA-Seq libraries were prepared and sequenced as described previously [88]. Briefly, total RNAs were extracted from lysates using a miRNeasy kit (QIAgen, #217004) following the manufacturer’s protocol for recovery of total RNAs (standard protocol). Genomic DNA contamination was depleted using Baseline-Zero DNase (Lucigen/Epicentre, #DB0715K) according to manufacturer’s instructions. Approximately 1 μg of DNase-treated RNAs were used to prepare a sequencing library. The rRNA depletion strategy using QIAseq FastSelect –rRNA Yeast Kit (Qiagen, #334215) was integrated into the RNA fragmentation step of the TruSeq Stranded mRNA Library Prep kit (Illumina, #20020594) according to the QIAseq FastSelect’s manual. Three libraries were multiplexed using recommended combinations of TruSeq RNA Single Indexes Set A (Illumina, #20020492) and sequenced (single-end, 75 cycles) on an Illumina NextSeq 500 sequencer.

### Sequence alignment

The yeast transcriptome used for sequence alignment was from https://github.com/Jacobson-Lab/yeast_transcriptome_v5, the generation of which was described previously [59]. Reads pre-processing, alignment, and quantification were performed on the University of Massachusetts Green High Performance Computing Cluster using the following provided software packages: cutadapt v1.9, bowtie v1.0.0, fastqc v0.10.1, samtools v0.1.19, bedtools v2.26.0, UMI-tools v1.1.1, and RSEM v1.3.0.

Ribosome profiling reads were pre-processed, aligned to the transcriptome, and transcript abundance quantified as described previously, except for the PCR duplicate removal step, where the UMI-tools software package was employed [89]. The UMI-tools’ “extract” function was used to record four nucleotides at each end of a read, which were introduced during library preparation by the NEXTflex Small RNA-Seq Kit v3. UMI-tools’ “dedup” function with the default (“directional”) method was used to identify and remove PCR duplicates based on the extracted UMIs. Number of reads processed, number of PCR duplicates removed, number of remaining unique reads, and other relevant sequencing statistics for each library are provided in S5 Table.

RNA-Seq reads were aligned to the transcriptome and transcript abundance quantified using RSEM without any pre-processing.

### Mass spectrometry (LC-MS/MS)

#### Sample preparation

Protein concentrations in cell lysates were determined by Pierce BCA Protein Assay according to the manufacturer’s protocol (Thermo Scientific). Aliquots of cell lysates containing 50 µg total protein were snap frozen, lyophilized in a SpeedVac, then reduced, alkylated, and digested following the S-Trap digestion protocol (ProtiFi). In brief, lyophilized lysates were first resuspended in 23 µl Lysis buffer (5% SDS in 50mM Triethyl ammonium bicarbonate (TEAB)). Resuspended protein extracts were reduced by adding 1 µl 200mM TCEP and incubating at 55 °C for 1 hour, then alkylated by adding 1 µl 375mM iodoacetamide (IAA) and incubating at room temperature for 30 minutes, protected from light. To further denature and trap proteins, samples were mixed with 2.5 µl of 27.5% phosphoric acid (H_2_PO_4_ in water) and 165 µl of binding/wash buffer (100mM TEAB in 90% methanol). The mixtures were applied to S-Trap columns and centrifuged at 4,000 g for 30 seconds. Columns were washed 5 times, each by 150 µl of binding/wash buffer and centrifugation at 4,000 g for 30 seconds. To digest proteins, 25 µl of digestion buffer containing 1 µg trypsin (5 µl of 0.2 µg/µl trypsin in 50mM TEAB + 20 µl 50mM TEAB) was added to each column and samples were incubated at 37 °C overnight. Digested peptides were collected by 3 subsequent centrifugations at 4,000 rpm for 1 min following the addition of these elution buffers for each collection: 1) 40 µl 50mM TEAB in water, 2) 40 µl 0.2% formic acid in water, and 3) 40 µl 50% acetonitrile in water. All flowthroughs from the same sample were pooled, lyophilized in a SpeedVac, and stored at -80 °C.

Samples were labeled using a TMT10plex labeling kit (Thermo Scientific). Lyophilized, digested peptides were resuspended in 50 µl 100mM TEAB and incubated with 10 µl of TMT10plex reagent (equilibrated to room temperature and resuspended in acetonitrile) at room temperature for 1 hour. Reactions were quenched with 5 µl of 5% hydroxylamine at room temperature for 15 minutes. Equal amounts (55 µl) of each sample were pooled together; 50 µl of the pooled reaction was saved for direct shotgun analysis and the rest for high-pH fractionation. For the latter, pooled samples were dried in a SpeedVac, resuspended in 300 µl of 0.1% trifluoroacetic acid (TFA) in water, and fractionated using a Pierce High pH Reversed-Phase Peptide Fractionation Kit (Thermo Scientific), collecting 1 flowthrough fraction, 1 wash fraction, and 6 step-gradient fractions.

#### Data acquisition

Mass spectrometry data was acquired using an Orbitrap Fusion Lumos Tribrid Mass Spectrometer (Thermo Scientific). Dried peptides were resuspended in 18 µl of 5% acetonitrile with 0.1% formic acid in water, vortexed for 2 minutes, and centrifuged at 16,000 rpm for 16 minutes. For mass spectrometry, 3.8 µl of the resuspended peptides were injected into the Mass Spectrometer. Peptides were trapped for 4 minutes at a flow rate of 4.0 µl/min onto a 100 µm I.D. fused-silica precolumn (Kasil frit) packed with 2 cm of 5 µm ReproSil-Pur 120 C18-AQ (dr-maisch.com), and eluted and separated in 120 minutes at a flow rate of 300 nl/min by an in-house made 75 µm I.D fused silica analytical column (gravity-pulled tip) packed with 25 cm of 3 µm ReproSil-Pur 120 C18-AQ (dr-maisch.com). Mobile phases were A (water (0.1% (v/v) formic acid) and B (acetonitrile (0.1% (v/v) formic acid). The biphasic elution program was as follows: 0-100 min (10-35% B); 100-120 min (35-65% B); 120-121 min (65-95% B); 121-126 min (95% B); 126-127 min (95-5% B); 145 min (STOP).

The MS data acquisition was performed in positive electrospray ionization mode (ESI+), within the mass range of 375-1500 Da with the Orbitrap resolution of 120,000 (*m/z* 200) and a maximum injection time of 50 milliseconds. Data dependent acquisition (ddMS2) was carried out with a 1.2 Da isolation window, a resolution of 30,000 (*m/z* 200), maximum injection time of 110 milliseconds, and the customed AGC target with a 38% of HCD collision energy.

#### Data analysis

Raw data files were processed with Proteome Discoverer (version 2.1.1.21, Thermo Scientific) and searched against the Uniprot Saccharomyces cerevisiae database (downloaded 06/28/2021) using Mascot Server (version 2.8, Matrix Science). Search parameters included full trypsin, with variable modifications of oxidized methionine, pyroglutamic acid (from Q), and N terminal acetylation. Fixed modifications were carbamidomethylation on cysteine and TMT10plex on peptide N-terminus and lysine side chain. Assignments were made using a 10ppm mass tolerance for the precursor and 0.05 Da mass tolerance for the fragments. Peptide and protein validation and annotation was done in Scaffold (version 5, Proteome Software, Inc.) using Peptide Prophet [90] and Protein Prophet [91] algorithms. Peptides were filtered at a 1% FDR, while protein identification threshold was set to greater than 99% probability and with a minimum of two identified peptides per protein. Protein clustering analysis was applied to increase the probability of protein identification for proteins that share peptides (e.g. paralogs). Quantitative analyses, including TMT label-based quantification, median normalization of log_2_ intensity values, and log_2_ fold change calculation were carried out in Scaffold Q+S.

Data acquired from flowthrough and wash fractions were used to determine the success of high-pH fractionation. Further analysis was based on data acquired from 6 step-gradient fractions.

### Bioinformatics and statistical analyses

Data analyses and visualization were performed in the R software environment versions 3.5 and 4.2 using the following R packages: data.table, dplyr, reshape2, readxl, openxlsx, caret, randomForest, rfPermute, rstatix, rcompanion, DESeq2, limma, Biostrings, seqinr, riboWaltz, ORFik, gprofiler2, scales, ggplot2, ggpubr, ggh4x, ggrepel, ggVennDiagram, ggseqlogo, patchwork, and Cairo.

#### Ribosome profiling analysis and readthrough efficiency calculation

Aligned reads of ribosome profiling libraries were processed by R package riboWaltz [92] for initial diagnostic, read length filter (retain reads 20-23 nt and 27-32 nt in length), and determination of read’s P-site offsets, which were manually checked and modified for accuracy (S6 Table). Read counts belonging to different mRNA regions (5’ UTR, CDS, extension, and distal 3’-UTR), read’s reading frame, and metagene analysis assessing periodicity were based on the read’s P-site location of mRNAs with annotated UTRs.

Readthrough efficiency was calculated for each mRNA as follows:

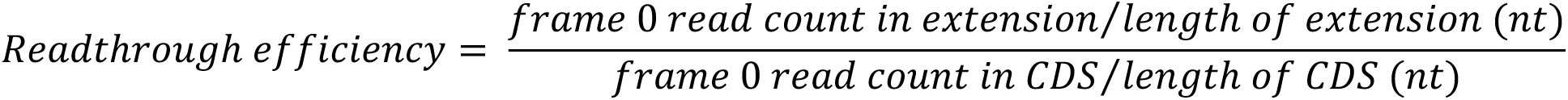

where the first 15 bp of the CDS region were excluded to avoid bias in ribosome accumulation over or near the start codon, and the extension region was defined as the 3’-UTR region from the canonical stop codon (inclusive) to the next in-frame stop codon (exclusive).

#### Random forest models

Random forest analyses were carried out with R packages caret [93], randomForest [94,95], and rfPermute [96]. For each sample, a random forest regression approach was trained to use mRNA features (previously defined in Mangkalaphiban et al. 2021 [59]) to predict readthrough efficiency values with 5-fold cross-validation, repeated 5 times, resulting in a total of 25 models. Each model was trained with 100 trees, the default number of features to split at each tree node (square root of number of features), and 1,000 permutation replicates to empirically determine p-value for feature importance. Feature importance score, percent increase in mean squared error (%IncMSE), was an average of scores extracted from 25 models. A feature was considered significantly important predictor of readthrough efficiency if its empirical p-value was less than 0.05 in at least 15 out of 25 models. Model performance metric was reported as an average of root mean squared error normalized to the range of readthrough efficiency values (NRMSE) across 25 regression models.

#### Analysis of transcript abundance changes

All analyses involving transcript abundance changes were performed with the R package DESeq2 [97]. The “expected_count” columns in the RSEM file output “isoforms.results” were used as input raw read count. Results were extracted with automatic independent filtering applied at significant cutoff (alpha) of 0.01. The false discovery rate (FDR) method was used to adjust the P-value.

For differential expression analysis of RNA-Seq libraries, mRNAs with adjusted P-value < 0.01 were considered significantly differentially expressed between samples, regardless of magnitude of log_2_ fold change. For Figures. 2C-D, S2C, and 6B, where RNA-Seq log_2_ fold change were plotted against mass spectrometry log_2_ fold change, expected_count of mRNAs in the same protein cluster were added together and RNA seq analysis was carried out as described.

For relative changes in translation efficiency (TE), ribosome profiling reads whose P-site locations were in the 5’-UTR, 3’-UTR, or the first 15 bp and the last 3 bp of the CDS (ribosomes paused over canonical start codon, translational ramp, and canonical stop codon) were discarded and transcript abundance for the rest of reads was re-quantified by RSEM. TE was defined as Ribo-Seq reads in CDS normalized to RNA-Seq reads. mRNAs with adjusted P-value < 0.05 were considered to have significant changes in TE between samples, regardless of the magnitude of log_2_ fold change.

#### Analysis of protein abundance changes

Differential abundance analyses of protein levels between samples were performed with the R package limma [98,99] on log_2_ normalized intensity data exported from Scaffold. The Benjamini-Hochberg method was used to adjust the P-value. Proteins with adjusted P-value < 0.015 were considered to have significant changes in abundance between samples, regardless of magnitude of log_2_ fold change.

Gene ontology analysis of proteins enriched (“Up”) or depleted (“Down”) was carried out by the R package gprofiler2’s gost function with default parameters [100,101].

#### mRNA features

mRNA features related to analysis of stop codon readthrough efficiency were defined as previously described [59].

Identification of uORFs in the 5’-UTR was done with the findORFs function in the R package ORFik [102], limiting uORF’s start codon to be AUG only and no minimum uORF length.

Poly(A) tracts in the 5’-UTR and 3’-UTR were defined as stretches of at least 10 consecutive adenines (the A in AUG of main ORF included), allowing at most 2 other nucleotides in the 10 A’s window.

Oligo(U) in the 5’-UTR and 3’-UTR was defined as a stretch of at least 7 consecutive uracils, allowing no other nucleotides in the window.

#### Motif discovery

Identification of sequence motifs enriched in 5’-UTR (excluding the start codon) or 3’-UTR (including the stop codon) sequences of mRNA TE groups compared to the Reference was carried out by the STREME algorithm from the MEME Suite 5.5.0 (https://meme-suite.org/meme/tools/streme) [75,103]. A general search for 3-15 nt-long motifs was performed as well as a focused search for shorter 3-6 nt-long motifs. Short motifs had to be identified in both searches to be regarded as significant enrichments.

#### Statistical analyses

Statistical analyses were performed using the R packages rstatix, rcompanion, and ggpubr. Specific statistical parameters, multiple-testing correction method, statistical significance (p-value or symbols representing ranges of p-values), and sample size (n) are reported accordingly on the figures, in the figure legends, or in Supplemental Table S2.

## Data Availability

Raw sequencing data have been deposited at the National Center for Biotechnology Information (NCBI) Gene Expression Omnibus (GEO) under accession numbers GSE229691 and GSE229692. The mass spectrometry proteomics data have been deposited to the ProteomeXchange Consortium via the PRIDE [104] partner repository with the dataset identifier PXD041495 and 10.6019/PXD041495. Analysis scripts and data used to generate figures are available at https://github.com/Jacobson-Lab/Pab1_deletion.

## ACKNOWLEDGMENTS

This work was supported by a grant to A.J. (1R35GM122468) from the U.S. National Institutes of Health. We thank Feng He, Chan Wu, Jill Moore, Zhiping Weng, Andrei Korostelev, and Sean Ryder for helpful discussions. We thank members of the UMass Chan Medical School Mass Spectrometry Core Facility for proteomics data acquisition, the UMass Chan Medical School Molecular Biology Core Labs for Fragment Analyzer service, and the UMass Chan Medical School Bioinformatics Core and Information Technology for the high performance computing platform.

## AUTHOR CONTRIBUTIONS

K.M. and A.J. conceived and designed the experiments, K.M. and R.G. carried out the experiments, K.M. wrote data processing scripts, K.M. and A.J. analyzed the data, K.M., and A.J. wrote the paper, and A.J. obtained funding for the study.

## SUPPORTING INFORMATION CAPTIONS

**S1 Fig. Replicate reproducibilities. A.** Correlation matrix showing Pearson correlation coefficients (*r*) of transcript abundance (non-zero RPKM values, log_10_-transformed) between pairs of sequencing libraries, RNA-seq and ribosome profiling libraries. **B.** Correlation matrix showing Pearson correlation coefficients (*r*) of log_2_ intensity values from mass spectrometry data between pairs of samples.

**S2 Fig. Changes in transcriptomes and proteomes of *pbp1Δ* and *pab1Δpbp1Δ* strains relative to WT. A.** Volcano plots of changes in transcriptome (RNA-Seq data) between strains. Orange, purple, and grey dots represent mRNAs with higher abundance (positive log_2_ fold change, adjusted p-value < 0.01), lower abundance (negative log_2_ fold change, adjusted p-value < 0.01), and no change (adjusted p-value ≥ 0.01), respectively. **B.** Volcano plots of changes in proteome (mass spectrometry data) between strains. Orange, purple, and grey dots represent proteins with higher abundance (positive log_2_ fold change, adjusted p-value < 0.015), lower abundance (negative log_2_ fold change, adjusted p-value < 0.015), and no change (adjusted p-value ≥ 0.015), respectively. **C.** Comparison of log_2_ fold change in transcriptome and proteome, with Spearman’s correlation coefficient. Grey, genes whose mRNA and protein abundance remained unchanged. Blue, genes whose protein but not mRNA abundance changed significantly. Red, genes whose mRNA but not protein abundance changed significantly. Green, genes whose mRNA and protein abundance both changed significantly.

**S3 Fig. Influences of mRNA features on readthrough efficiency. A.** Average ± standard deviation of performance metrics (normalized root mean squared error (NRMSE)) extracted from 25 random forest models (5-fold cross-validation, repeated 5 times) trained for each strain to predict readthrough efficiency. **B**-**C**. Heatmaps of median readthrough efficiency of mRNA groups, grouped by the identity of the stop codon or the identity of nucleotide at positions near the stop codon (**B**) or the identity of P-site codon (**C**), relative to median readthrough efficiency of all mRNAs in the sample. Positive (red) and negative (blue) values indicate that the group has higher and lower readthrough efficiency than the sample median, respectively. Two-sided Wilcoxon’s rank sum test with Benjamini-Hochberg method for multiple-testing correction was used to compare a group’s median readthrough efficiency to the sample median. Significant results (p < 0.05) are represented as bigger tiles.

**S4 Fig. Influences of start codon context in changes of translation efficiency (TE). A-B.** Relative proportions of nucleotide usage upstream (**A**) and downstream (**B**) of main ORF’s AUG (positions +1 +2 +3) in Up and Down groups relative to Reference. **C.** Relative proportions of nucleotide usage from the mRNA 5’ cap (first 18 nucleotides of the 5’-UTR sequences) in Up and Down groups relative to Reference. In all panels, Reference (Ref.) group includes all mRNAs regardless of TE changes (Up + Down + Unchanged) to recapitulate the general proportions in the transcriptome. Positive (red) and negative (blue) log_2_ relative proportion indicates that the nucleotide is over represented and under-represented, respectively, in the group compared to the Reference. Pairwise χ^2^ test with Benjamini-Hochberg method for multiple-testing correction was used to compare the nucleotide frequencies between Reference, Up, and Down groups. p < 0.05 for Up or Down vs. Reference is represented by a big tile, while non-significant results are represented by a small Tile. p < 0.05 for Up vs. Down is represented by an asterisk (“*”).

**S1 Table. Gene ontology analysis results of proteins with significant changes in abundance in *pab1Δpbp1Δ* cells relative to *pbp1Δ* cells.**

**S2 Table. Statistical test results of pairwise comparison of mRNA features between Reference, Up, and Down TE groups.** Related to Figs 4B, 4D, 4E, 4F, 5D, and 5F.

**S3 Table. List of yeast strains and genotypes used in this study.**

**S4 Table. List of oligonucleotides used in this study.**

**S5 Table. Ribosome profiling reads processing and alignment statistics.**

**S6 Table. Ribosome footprint P-site offsets from footprint’s 5’ end and 3’ end**

**S1 File. Interactive visualization of gene ontology results from S1 Table.** Interactive Manhattan plot was generated by the gostplot function in the R package gprofiler2 [101]. Each circle on the plot represents a gene ontology (GO) term. The size of the circle reflects the number of genes in the GO term. GO term name, ID, size, and adjusted p-value can be seen by hovering over the circle. GO terms are grouped and colored by data sources on the x-axis. GO terms that are closer in hierarchy are also closer visually. The y-axis shows adjusted p-values in negative log10 scale. The plot is capped at 16, meaning GO terms with adjusted p-value < 10^-16^ are plotted at “>16” on the y-axis.

