## Supplementary figures and images for "Direct and indirect consequences of *PAB1* deletion in the regulation of translation initiation, translation termination, and mRNA decay"

### Supplemental Figs. 1-4

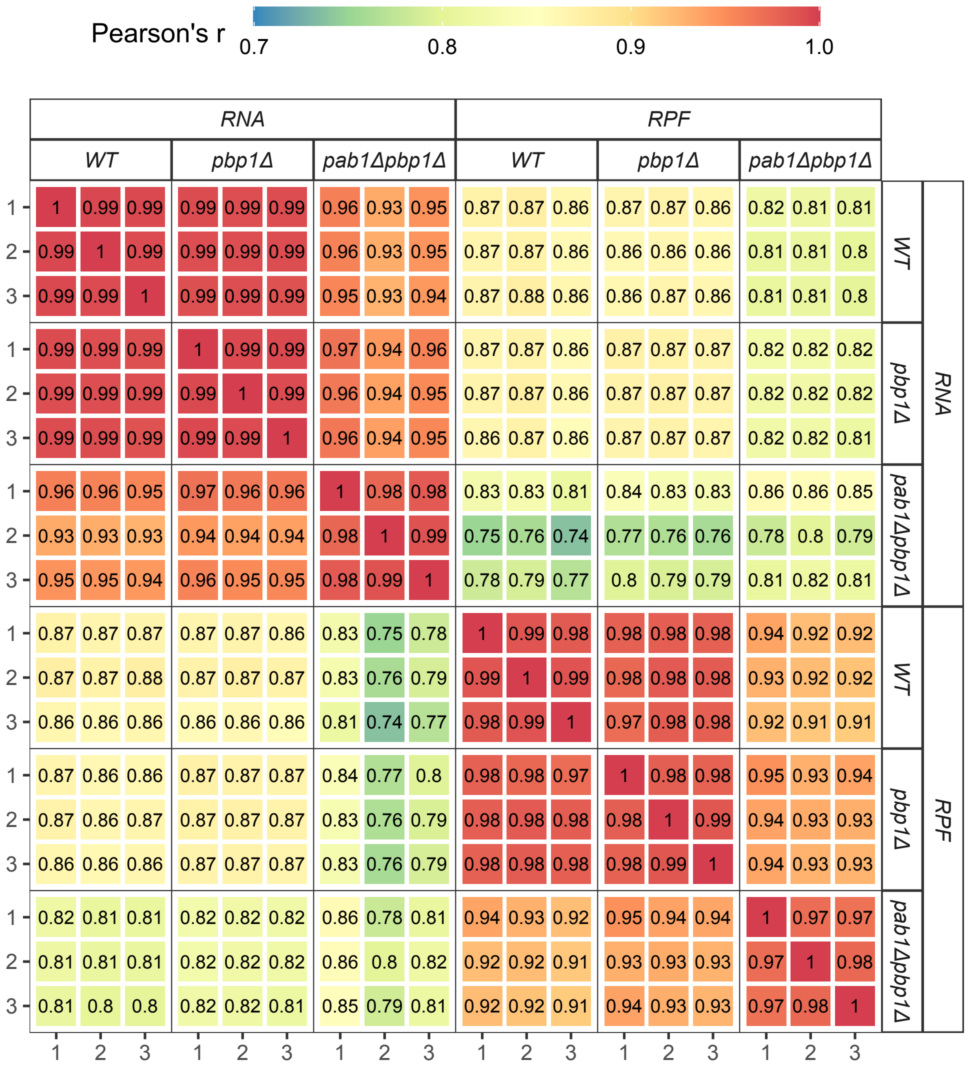

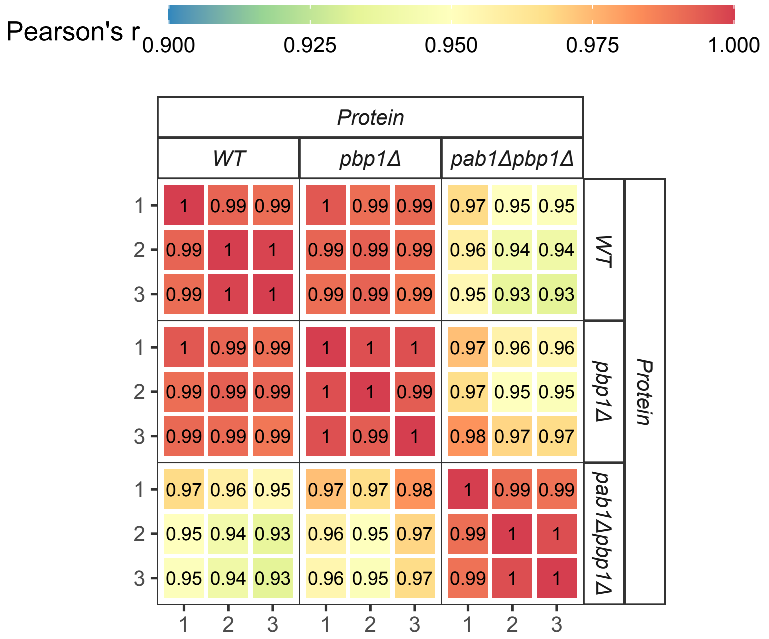

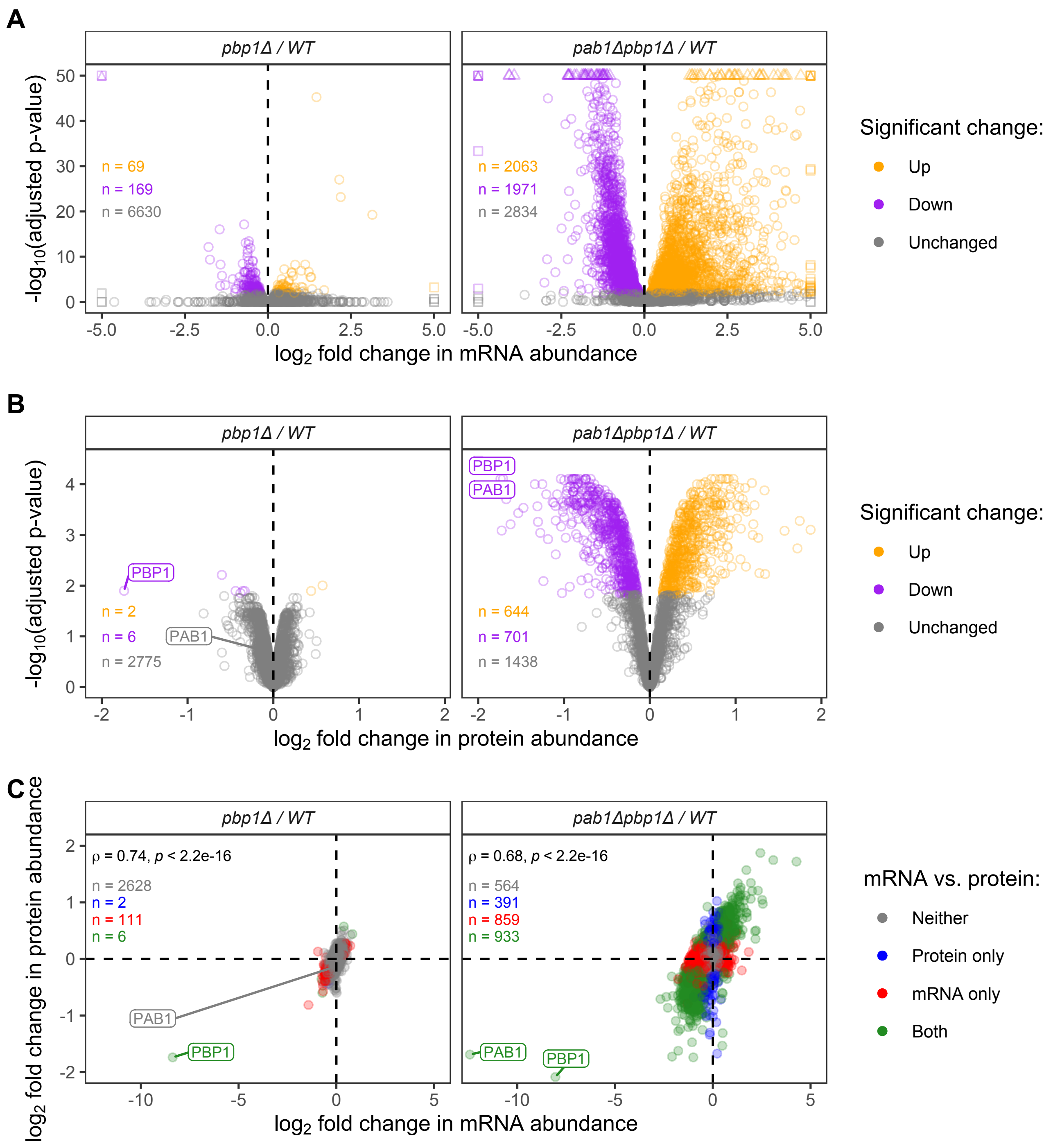

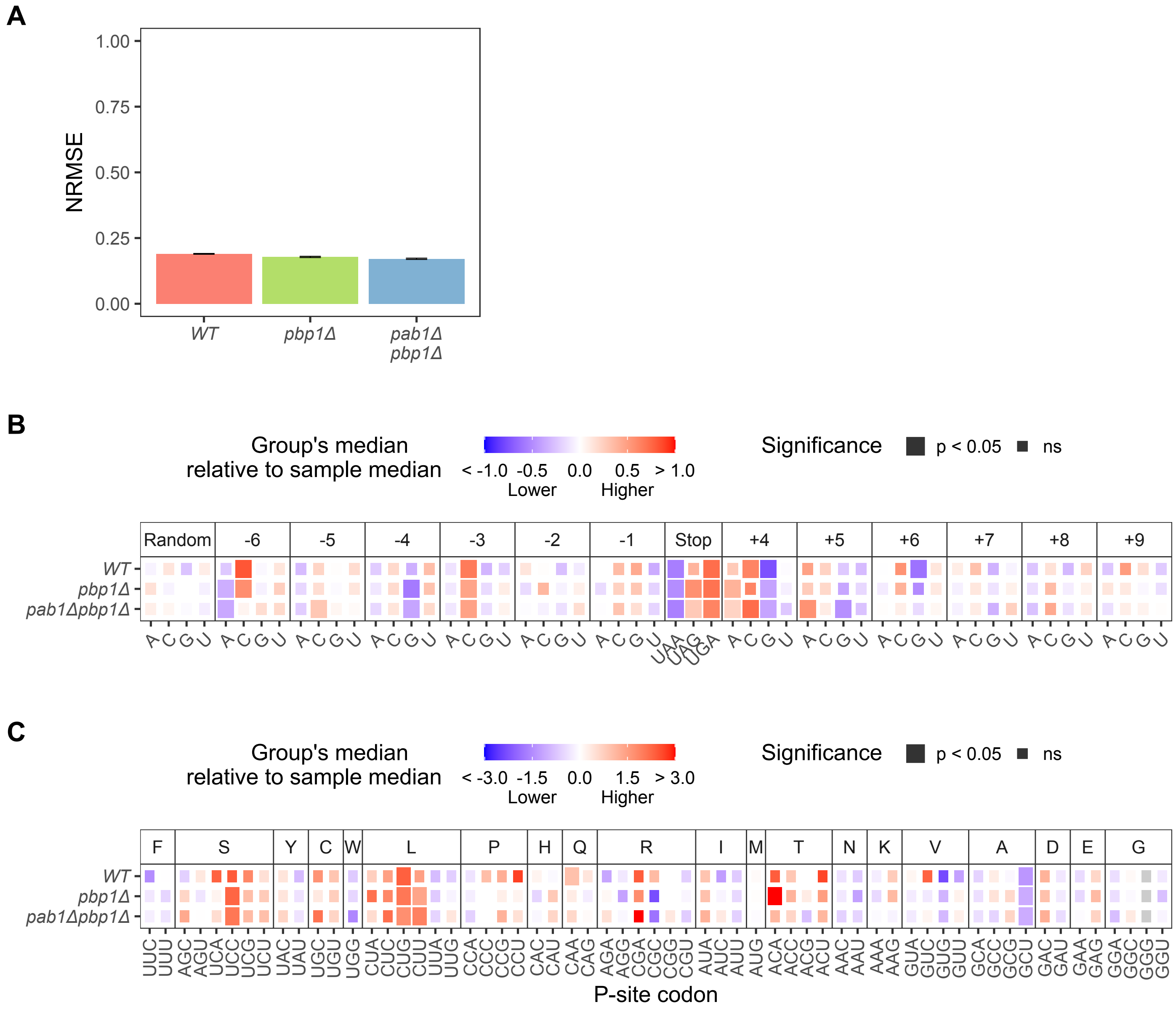

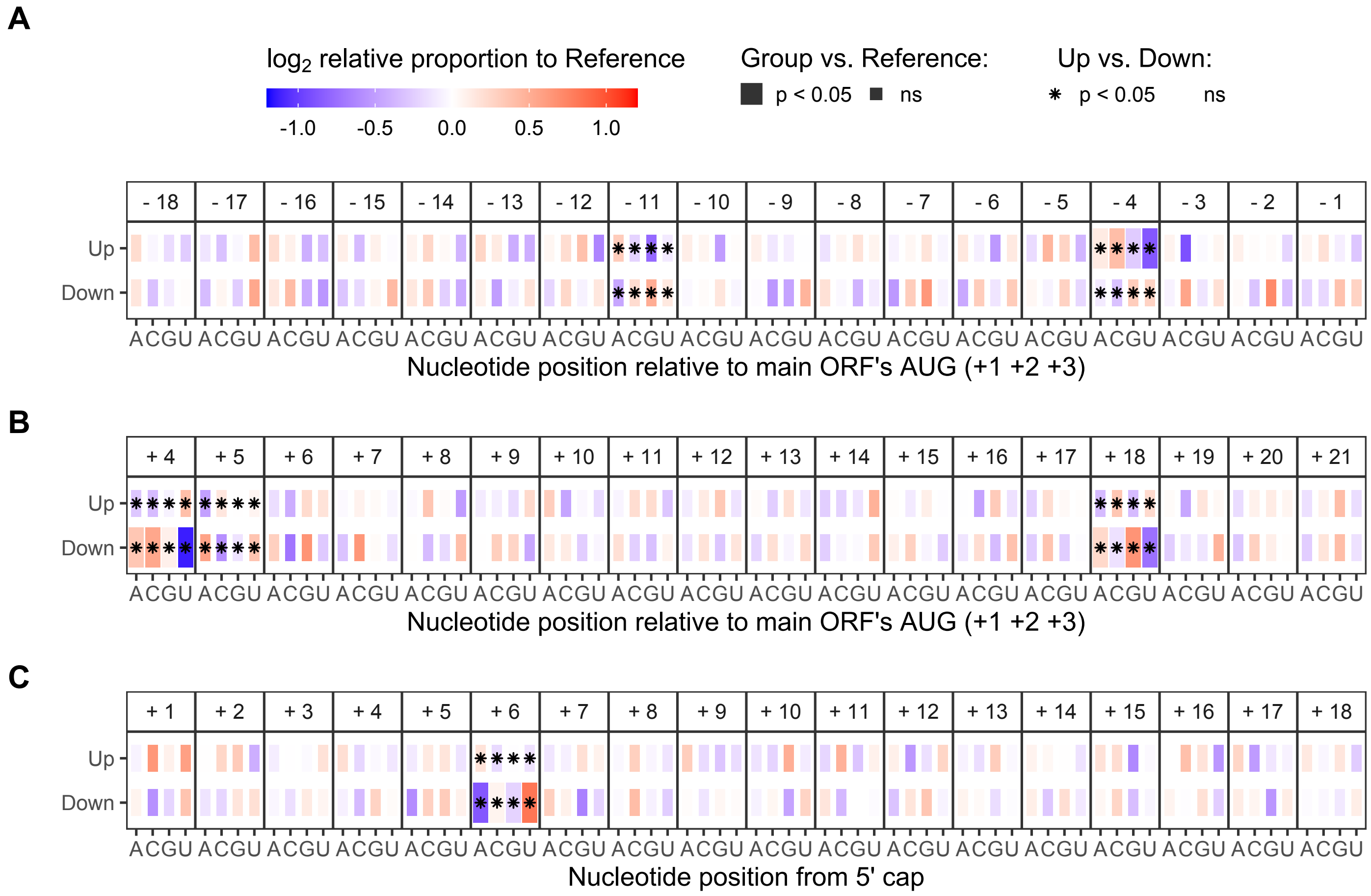
